# Lysosomal multi-omics reveals altered sphingolipid catabolism as driver of lysosomal dysfunction in the aging brain

**DOI:** 10.1101/2025.09.10.675421

**Authors:** Chinmoy Sarkar, Yi Chen, Dexter PH Nguyen, Mehari M Weldemariam, Yulemni Morel, Amir Mehrabani-Tabari, Nikki Gorny, Olivia Pettyjohn-Robin, Nivedita Hegdekar, Sagarina Thapa, Sazia Arefin Kachi, Sabrina Bustos, Stephanie Zalesak-Kravec, Christina Williams, Nicholas Leahy, Renee Ti Chou, Shilpa D Kumar, Carrie McCracken, Thomas A Blanpied, Mariusz Karbowski, Jace W Jones, Maureen A Kane, Michael P Cummings, Marta M. Lipinski

## Abstract

Recent data indicate that lipid composition has profound influence on the brain function and that changes in lipid homeostasis affect brain aging and predisposition to neurodegenerative diseases. Lipids dynamically reside in multiple intracellular locations and their organellar distribution is important for specific interactions and biological function. During brain aging lipid changes have been specifically noted in lysosomes, but the identity of the accumulated lipids, their interactions with other biomolecules such as proteins, and their functional relevance have not been characterized. We used mass spectrometry (MS) to assess longitudinal changes in the lipidome and proteome of lysosomes isolated from the mouse cortex, from the age of 3- to 24-months. Our statistical and machine learning analyses identified two factors demonstrating predictive power for age and differences in both lipids and proteins. Of these, factor 1 was the best predictor of sample age. Factor 1 lipids with the highest feature importance included multiple species of hexosylceramides (HexCer) and their sulfonated derivatives, sulfatides (SHexCer), all of which increased with age. Increased factor 1 proteins included myelin proteins, select sphingolipid catabolism enzymes and proteins associated with lysosomal storage diseases. Our analyses suggested that mechanisms underlying factor 1 encompass the combination of an age-dependent increase in lysosomal delivery of myelin components and alterations in lysosomal sphingolipid catabolism favoring degradation of sphingomyelin over HexCer. The overall age-related lysosomal changes resembled those observed in lysosomal storage diseases, particularly Gaucher disease, where accumulation of HexCer species is associated with lysosomal dysfunction. To corroborate factor 1 predictions, we employed a combination of biochemical, imaging and flow cytometry approaches, which confirmed alterations in sphingolipid catabolism and lysosomal accumulation of myelin components. These changes were associated with age-related alteration in lysosomal morphology, lysosomal dysfunction and inhibition of autophagy in both neurons and microglia. Our findings indicate that factors contributing to lysosomal aging resemble those observed in lysosomal storage diseases and underscore the significance of organelle-specific analyses for dissecting mechanisms contributing to brain aging.

## Introduction

Because of the prevalence of branched cells with high membrane/volume ratio and the presence of lipid-rich myelin^1,2^ brain has higher level of lipids than any other tissue except the adipose^3,4^. Importance of lipid for brain structure and function is further reflected in its unique lipid composition, and the exceptional diversity of lipid species present in the neural tissue^5^. Lipids are major components of cellular and organellar membranes, affecting their structure, fluidity and barrier function^6,7^. Many lipids also play important roles in cellular signaling, either by themselves functioning as signaling intermediates, or by impacting the formation and function of lipid rafts which serve as docking platforms for intracellular signaling molecules^8^. Consistent with their importance for brain function, alterations in lipid homeostasis are associated with the brain aging process and with increased susceptibility to neurodegenerative diseases^9^. For example, polymorphisms in genes involved in lipid handling, such as *APOE* and *TREM2* are implicated in late onset Alzheimer’s disease (AD)^10,11^, while altered expression of lipid metabolism genes is common in age- and disease-associated microglial populations^12–14^.

Lipids dynamically reside in multiple intracellular locations and their organellar distribution is important for specific interactions and biological function^15^. Distinct organellar compartments establish specialized local environments that facilitate interactions between lipids and other macromolecules, such as proteins, thereby enabling physiological processes. Consequently, organellar localization must be considered when determining the significance of alterations in lipid abundance and function^15–17^. Cellular lipid metabolism is also highly compartmentalized, with lipid synthesis carried out mainly in the endoplasmic reticulum (ER) and peroxisomes, and degradation occurring in mitochondria, peroxisomes and lysosomes. Specific lipid accumulation in the endo-lysosomal compartments has been noted in the aged and neurodegenerative disease brain^18^. However, neither the identity of accumulating lysosomal lipids, their interactions with other biomolecules residing in the same cellular compartment, nor their specific effects on organellar, cellular and overall brain function has been characterized.

Lysosomes are essential centers of cellular catabolism, playing a crucial role in the breakdown of proteins, lipids and organelles. Age-related neurodegenerative diseases are characterized by accumulation of protein aggregates such as amyloid β, tau and α synuclein, and of lipid storage materials such as lipofuscin, indicating defects in cellular degradation pathways^19^. Defects in autophagy, a lysosome-dependent catabolic pathway, are observed in multiple neurodegenerative diseases, including Alzheimer’s (AD) and Parkinson’s disease (PD), and thought to contribute to accumulation of pathological protein aggregates and defective organelles^20,21^. Autophagy also affects inflammatory responses, with high levels of autophagy associated with anti-inflammatory, and low levels with pro-inflammatory phenotypes^22,23^. Decline in autophagy-lysosomal function has been observed during normal brain aging but the contributing mechanisms are not fully understood^18,24^.

Sphingolipids are a group of bioactive lipids including ceramides (Cer), hexosylceramides (HexCer) and sphingomyelins (SM), which are characterized by the presence of lipid moiety on an aliphatic base known as sphingosine^25^. Accumulation of sphingolipids is commonly observed in lysosomal storage diseases (LSDs) where loss-of-function mutations in lysosomal lipid catabolism enzymes cause their respective substrates to accumulate. This results in impairment of both degradative and signaling functions of the lysosomes and a broad range of cellular defects, including loss of proteostasis, disruption of autophagy and endocytic trafficking^25–28^. Parallels between the etiology of LSDs and neurodegenerative diseases, especially PD, frontotemporal dementia and amyotrophic lateral sclerosis (ALS), have been noted^29,30^. Changes in sphingolipids have been also reported in AD; however, their significance remains poorly understood^31^.

Here we elucidate the role of lysosomal lipids during brain aging by analyzing lipid and protein composition of lysosomes isolated from murine cortex, from 3- to 24-months of age. Our analyses implicate increase in lysosomal delivery of myelin components and alteration in lysosomal sphingolipid catabolism favoring degradation of sphingomyelin over HexCer as drivers of age-related lysosomal dysfunction. The overall changes resemble those observed in lysosomal storage diseases, particularly Gaucher disease. We employ a combination of experimental approaches to corroborate the multi-omics results and demonstrate that altered lysosomal lipid composition is associated with changes in lysosomal morphology, lysosomal dysfunction and inhibition of autophagy in both neurons and microglia. Our findings identify lipid-dependent mechanisms contributing to lysosomal and autophagy dysfunction during brain aging and underscore the significance of organelle-specific analyses for understanding lipid function.

## Results

### Purification and characterization of mouse cortical lysosomes

We used density gradient centrifugation to prepare lysosome-enriched fractions from the mouse cortex for mass spectrometry (MS)-based lipid and protein analyses (**Figure 1a**)^16^. The lysosome-enriched fractions showed high abundance of lysosomal proteins, especially their mature proteolytically processed forms associated with lysosomal maturation (**Figure 1b**). Conversely, markers of other cellular compartments including mitochondria, endosomes and ER were either decreased or unaltered. Additionally, we detected 2.8-fold increase in the activity of lysosomal protease cathepsin D (CTSD) as compared to whole brain lysates (**Figure 1c**). LysoTracker-Red staining confirmed that approximately 89% of purified vesicles retained membrane integrity and acidic pH (**Figure 1d**). These data suggest that the lysosome-enriched fractions consisted primarily of acidic vesicles including lysosomes, late endosomes and autolysosomes.

**Figure 1.**
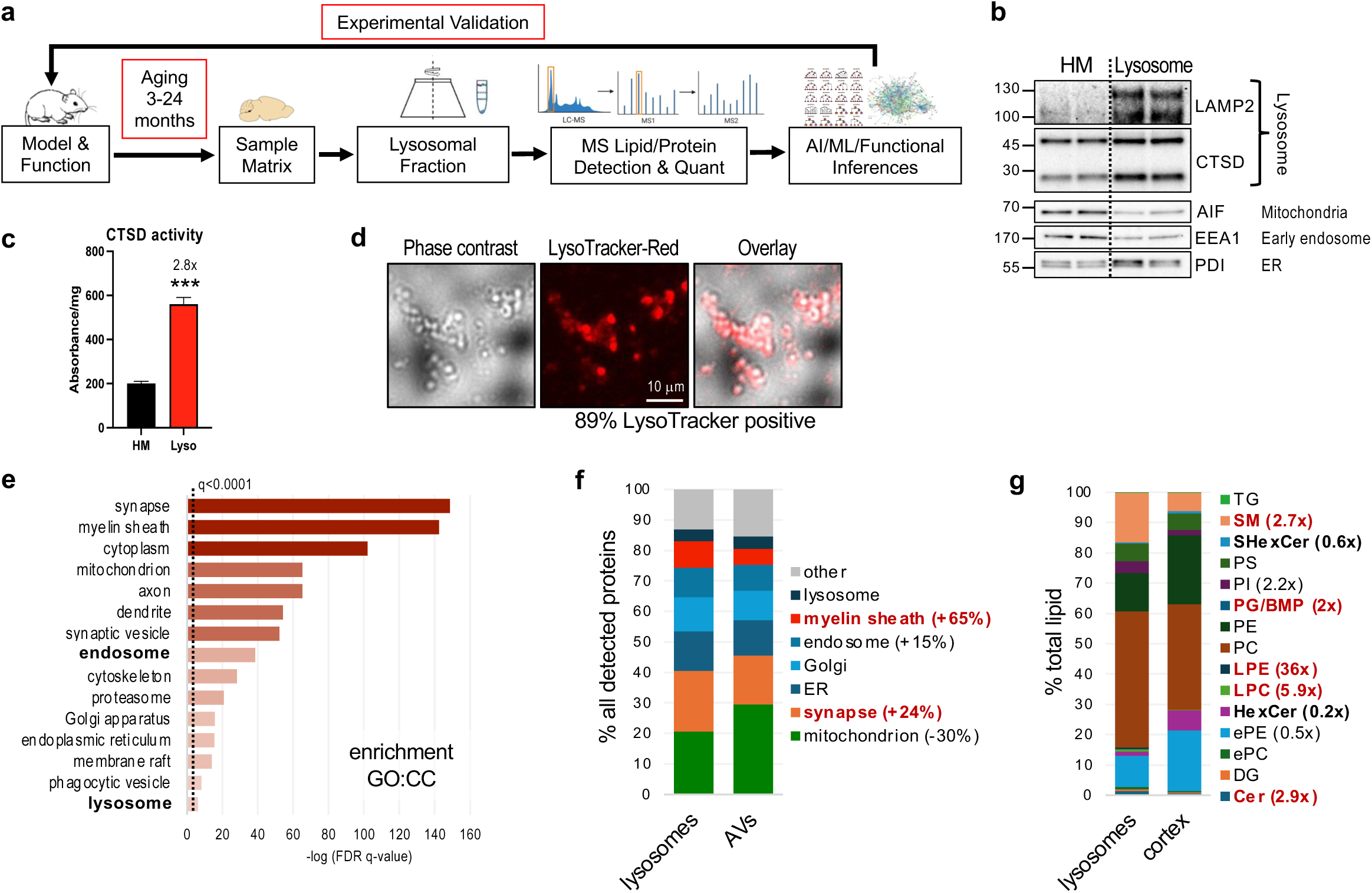
Preparation and characterization of mouse brain lysosomes. **a,** Outline of the overall experimental design. Mice were subject to aging and characterization, followed by collection of cortical tissues and preparation of lysosome-enriched fractions between 3 and 24 months of age. Lysosomes were subject to mass spectrometry (MS) lipid and protein detection and quantification, followed by machine learning (ML) based data integration and interpretation. ML functional inferences were used as basis for experimental validation. **b-d,** Validation of lysosomal-enrichment procedures, including Western blot demonstrating enrichment for lysosomal as compared to other compartment markers in lysosomal fractions vs total brain homogenates (HM) (**b**); comparison of cathepsin D (CTSD) activity in purified lysosomal fractions versus brain homogenates (**c**); and lysotracker staining demonstrating that approximately 89% of purified vesicles are acidic (**d**). **e,** Gene Ontology Cellular Compartment (GO:CC) analyses of all proteins detected in lysosomal fractions from 3-month-old mouse cortex. Mouse genome was used as background. **f**, Comparison of protein composition between lysosomes versus autophagosomes purified from young mouse brain (Kallergi et al., 2023, Neuron 111:2329–47) based on GO:CC analyses. **g,** Comparison of lipid classes detected in 3-month-old mouse cortex lysosomes versus total cortical lysates.

Lysosome-enriched fractions were subjected to parallel lipidomic and proteomic analyses using established liquid chromatography-tandem mass spectrometry workflows. Total lipids were analyzed by liquid chromatography coupled to high-resolution tandem mass spectrometry^32–34^. The same samples were in parallel subjected to label-free quantitative proteomic analyses using nano-flow liquid chromatography coupled to high-resolution tandem mass spectrometry^35–37^. In 3-month-old lysosomal samples we identified an average of 2,636 proteins and ∼500 lipids per sample (**Supplementary Tables S1 and S2**). As expected, we detected numerous lysosomal (ex. LAMP1/2, CTSB/D/F, PSAP, MTOR; 76 proteins total), autophagosomal (ATG9A, GABARAPL1/2, WDFY3/ALFY; 15 total) and endocytic pathway (RAB7, RAB5A/B/C, SORT1, SNX3; 185 total) proteins. Other enriched classes based on Gene Ontology Cellular Component analysis (GO:CC), included mitochondrial (395, 21%), synaptic (379 total, 20%) and ER (245, 13%) proteins (**Figure 1e, Supplementary Figure S1a**). This is consistent with previous data indicating enrichment of proteins derived from these compartments in purified mouse brain autophagosomes, which constitute one of the major lysosomal cargo delivery routes (**Figure 1f, Supplementary Figure S1b**)^38,39^. The mitochondrial proteins included both inner and outer membrane proteins, respiratory chain, matrix and mitochondrial DNA-associated proteins as well as several mitophagy receptors (NIPSNAP1, FKBP8, PHB2). Consistent with previous analyses of brain-derived autophagosomes, neither PINK1 nor parkin (PRKN) were present^38^. We did detect other selective autophagy receptors, including ER-phagy (RTN3, ATL3) and aggrephagy (ALFY, TOLLOP)^39^. Other enriched proteins included cytoskeleton-associated proteins (255, 13%), consistent with association with cytoskeleton necessary for lysosomal and other vesicle movement and intracellular localization, and myelin components (164, 8.6%). As compared to the reported autophagosomal contents, we noted increased proportion of synaptic (23% increase) and myelin sheath (65% increase) and decrease in mitochondrial (30% decrease) proteins^39^. We expect these differences may reflect extracellular endocytic/phagocytic in addition to intracellular autophagic cargo delivery to the lysosomes.

Lipid species detected in cortical lysosomes from 3-month-old mice belonged to multiple lipid classes (**Supplementary Table S2**). We compared distribution of lysosomal lipid classes to those observed in whole cortical tissue at the same age (**Figure 1g**). Although our overall lipid class distribution in cortical tissue was comparable to previous reports^4^, we detected significant changes in lipid class representation between cortical tissue and lysosome-enriched fraction. Most significant was enrichment in lysophospholipids (36-fold for lyso-phosphatidylethanolamine, LPE, and 5.9-fold for lyso-phosphatidylcholine, LPC). These increases are consistent with the role of lysosomes as hubs for catabolism of membrane-derived phospholipids, which are degraded by phospholipase families A and B (PLA and PLB) to generate lysophospholipids^40^. Consistent with our proteomic data showing enrichment in myelin components, we also detected increase in sphingomyelins (2.7-fold) and their degradation products, ceramides (2.9-fold). Conversely, levels of hexosylceramides (HexCer) and their sulfonated derivatives sulfatides (SHexCer) were decreased (0.2-fold for HexCer, 0.6-fold for sulfatides). Similarly to sphingomyelins, HexCer and sulfatides are abundant in myelin and catabolized in lysosomes to generate ceramides; their low abundance may reflect efficient degradation in young animals. Finally, we observed higher abundance of bis(monoacylglycero)phosphate (BMP, also known as lysobisphosphatidic acid, LBPA), a lipid species rare in most organelles and compartments of mammalian cells but enriched in lysosomes, where it plays a role as anchor and co-factor for catabolic enzymes^41^.

### Multi-omics of cortical lysosomes as a function of aging

To determine lysosomal lipidome and proteome changes occurring during brain aging, we purified lysosome-enriched fractions from cortices of 12-month-old (middle aged), 18-month-old and 24-month-old (aged) mice and from 3-month-old (young) controls. To ascertain that observed lysosomal changes correlate with changes in brain function, prior to harvest mice were subject to longitudinal behavioral assessments, including both motor (beam walk) and cognitive (Morris Water Maze and novel object recognition) function. As expected, all assessments confirmed progressive decline in function, which became significant at 18 months and further exacerbated at 24 months as compared to young mice (**Supplementary Figure S2**).

Lysosomal fractions from all age groups plus corresponding young controls were subjected to parallel MS-based lipidomic and proteomic analyses. We incorporated several study design elements to control for batch effects inherent in longitudinal omics studies. This included groups of young control lysosomal samples purified at the same time as the corresponding aged samples, as well as identical batches of young lysosomes derived from pooled control samples included with each of the age groups. This approach allowed us to identify sources of any observed batch effects (sample collection versus MS instrument run) and to apply robust computational methods for batch correction.

Overall, 2986, 2004, and 2287 proteins were quantified at 12, 18, and 24 months, respectively. Of these, 1554 proteins were consistently detected in all samples across the three age groups. Approximately ∼500 lipids were detected in all age groups. However, approximately half of the lipids had low ID confidence or low/variable abundance and were excluded from further analyses. This resulted in a list of 234 high confidence distinct lipids that were present in all age groups

### Age-dependent changes in brain lysosomal lipidome

We examined changes in lysosomal lipidome during brain aging. Pairwise principal component analyses (PCA) demonstrated that aging was associated with significant changes in lysosomal lipid abundance as compared to lysosomes from 3-month-old control mice (**Figure 2a-c**). Overall, at 12 months we detected 151 differentially abundant lipids, 157 at 18 months and 91 at 24 months (*P* < 0.05, FDR 5%, **Supplementary Tables S3-S8**). Over time larger proportions of lipids were increased rather than decreased in the aged lysosomes (53% increased at 12 months, 56% at 18 months, 64% at 24 months, **Figure 2d-f**). Since many lipids are degraded in the lysosomes, this suggests age-related decline in the ability of lysosomes to catabolize lipids. This is consistent with previous reports of accumulation of lipofuscin and other lipid byproducts in the endo-lysosomal compartments of the aging brain^18^.

**Figure 2.**
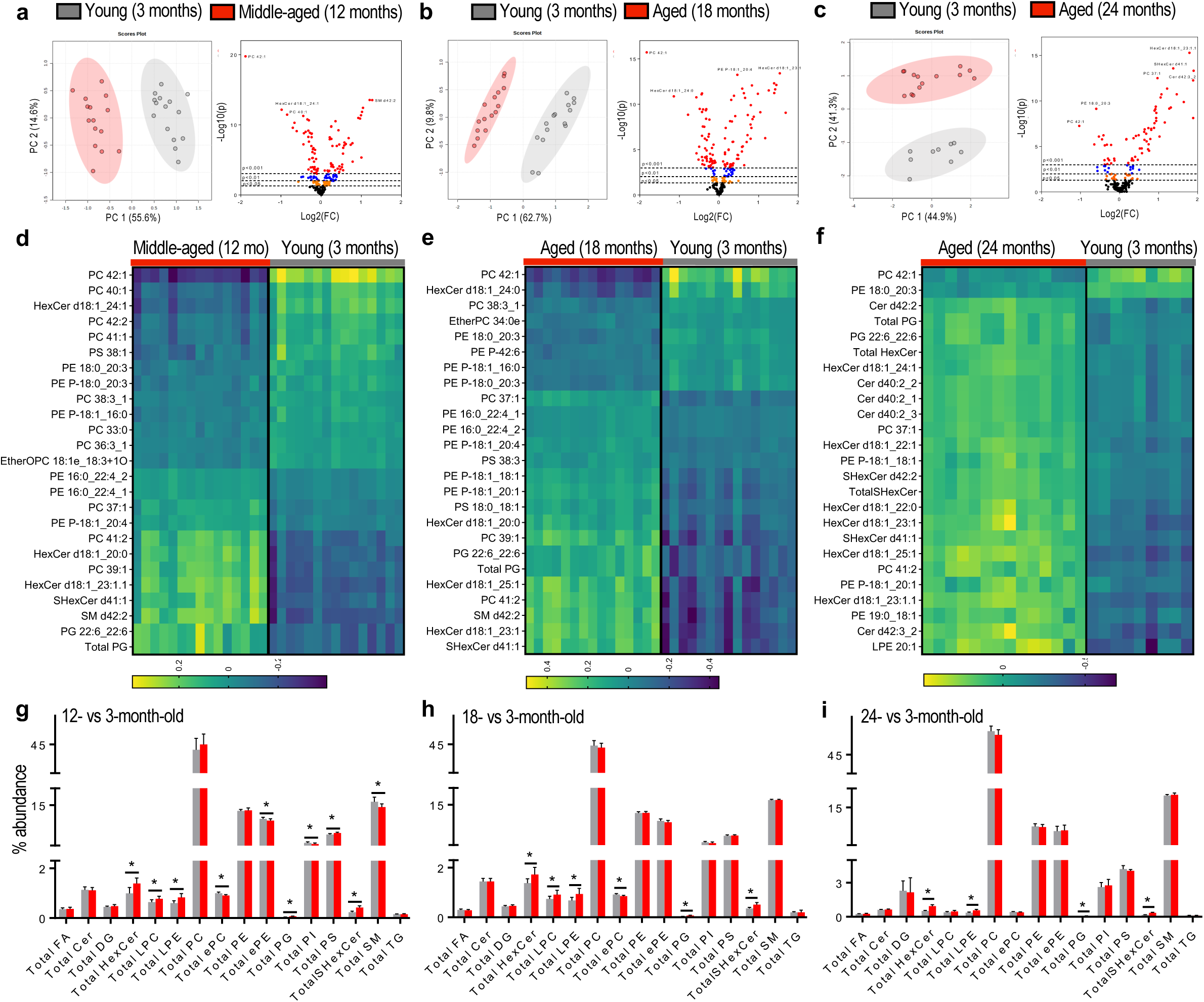
Lipidomic changes in cortical lysosomes as a function of age. **a-c,** Principal component analysis (PCA) and volcano plots for pairwise comparisons between lipid composition of lysosomes from control (3-months-old) and aged mouse cortices. PCA plots demonstrate clear group separation, while volcano plots highlight a substantial number of differentially abundant lipids. **d-f,** Hierarchical clustering (top 25 lipids) reveals distinct stratification between aged and control samples. Differentially expressed lipids were used as input for visualization using MetaboAnalyst (*version*, *5.0*). **g-i,** Lipid class abundance analyses comparing aged to corresponding young cohorts. Data represent sum composition of individual lipids per lipid class (SI Table S3). Bar graphs are mean ± SD; *p-value < 0.05; student t-test.

Aging altered abundance of several different classes of lysosomal lipids. This included age-dependent increase in lysophospholipids (including LPC and LPE, **Figure 2g-i**) which could reflect either age-dependent increase in phospholipid processing by lysosomal phospholipases or inability of aged lysosomes to export resulting lysophospholipids. The unique structure of lysophospholipids (one rather than two lipid moieties attached to the polar head) confers upon them detergent-like properties and their accumulation can cause membrane damage and permeabilization^42,43^. Cleavage of membrane phospholipids to lysosphospholipids also leads to release of free fatty acids, which can serve as precursors for generation of inflammatory modulators such as leukotrienes and prostaglandins, suggesting potential contribution to age-related neuroinflammatory changes. At 12 and 18 months we also detected decrease in ether lipids (ePC and ePE). Double bonds present in ether lipids can serve as sacrificial antioxidants and their decline suggests greater susceptibility to and/or presence of oxidative damage^43^.

The most consistently altered group of lysosomal lipids were sphingolipids. This included pronounced increase in hexosylceramides (HexCer, including glucosyl and galactosyl species) and sulfatides (SHexCer). Changes in individual sphingolipid species were apparent in all age groups but were exacerbated over time, with 20% of the 25 most altered lipids belonging to this class at 12 months, 24% at 18 months and 60% at 24 months (**Figure 2d-f**). In addition to progressive accumulation of HexCer and sulfatides, we observed discontinuous changes in levels of sphingomyelins (SM), including overall decrease in SM abundance at 18 but not 12 or 24 months (**Figure 2h**). Another class of lipids accumulating with age were phosphatidylglycerols (PGs) and/or their structural isomers, BMPs. BMPs are rare in most cellular compartments but enriched in lysosomes and late endosomes where they play a role in regulation of sphingolipid catabolism^41^.

### Age-dependent changes in brain lysosomal proteome

Pairwise comparisons between aged and young samples reveled significant changes in lysosomal proteomes at 12, 18 and 24 months as compared to 3-month-old controls (**Figure 3a-c**). Similarly to lipids, the proportion of proteins with higher lysosomal abundance increased over time, with 52%, 64% and 88% of the top 25 most altered proteins accumulating at 12, 18 and 24 months, respectively (**Figure 3d-f, Supplementary Tables S9-S14**). To determine whether observed proteomic changes may reflect age-dependent changes in gene expression, we performed bulk RNA sequencing analyses on mouse cortical tissues at 3, 12 and 18 months old (**Supplementary Table S15-S17**). Our data demonstrate an overall lack of significant correlation between age-dependent changes in lysosomal protein abundance and cortical gene expression (**Supplemental Figure S3a-b**). This suggests that most lysosomal proteome changes do not reflect altered gene expression but rather are driven by post-transcriptional factors such as protein translation, localization and stability. This is consistent with previous studies demonstrating poor correspondence between transcriptomic and proteomic data^44–46^, and supports the need for organelle-specific evaluation to understand lipid and protein function and interactions.

**Figure 3.**
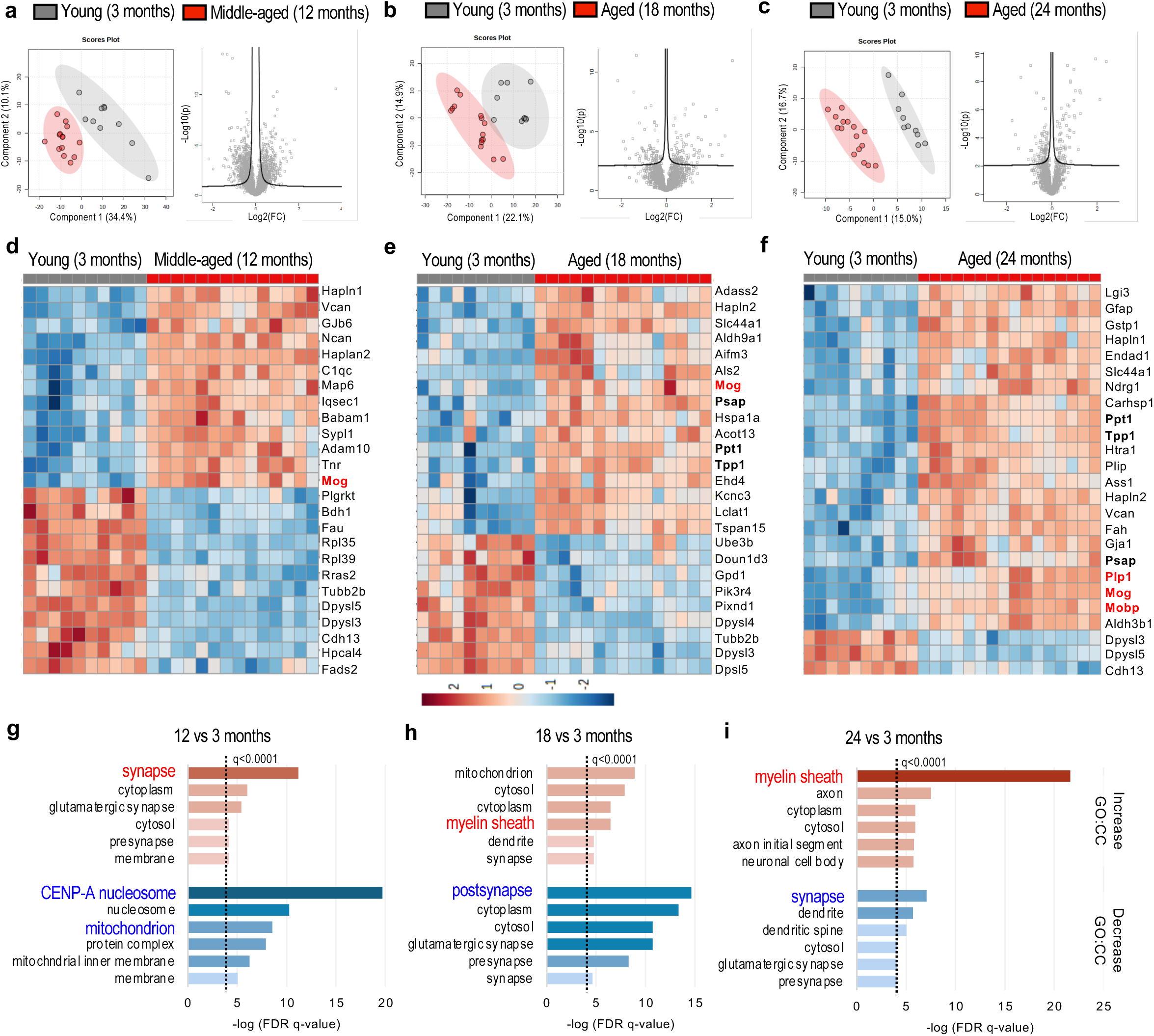
Proteomic expression changes in cortical lysosomes as a function of age. **a-d,** Principal component analysis (PCA) and volcano plots for pairwise comparisons between protein composition of lysosomes from control (3-months-old) and aged mouse cortices. PCA plots demonstrate clear group separation, while volcano plots highlight a substantial number of differentially abundant proteins. **d-f,** Hierarchical clustering (top 25 proteins) reveals distinct stratification between aged and control samples. Differentially expressed proteins were used as input for visualization using MetaboAnalyst (*version*, *5.0*). **g-h,** Gene Ontology Cellular Compartment (GO-CC) analyses (NIH DAVID, v2025_1) of proteins with significantly (q<0.05) altered (red – increased, blue – decreased) abundance as compared to young controls. All proteins detected in lysosomal preparations at each age were used as background.

We used Gene Ontology Cellular Component (GO:CC) analysis to determine sources of proteins altered at each age. Consistent with data indicating decrease in mitophagy and increase in synapse-derived autophagosomal cargos in mature versus adolescent mouse brain^39^, at 12 months the most significantly enriched proteins were those associated with synapses, while mitochondrial proteins were decreased (**Figure 3g**). However, the most decreased class at this age were nucleosomal proteins. This possibly reflects decrease in overall cell proliferation during brain maturation as well as potential decline in homeostatic nucleophagy in addition to mitophagy^47,48^. Unlike the 12-month group, at 18 and 24 months synaptic and other neuronal projection derived proteins were observed among both the increased and the decreased groups. Additionally, while the overall enrichment in neuronal and synaptic proteins in aged lysosomes resembled what was previously observed in aged autophagosomes^39^, we noted only few overlaps between specific altered protein cargoes (**Supplementary Figure S3c-e**).

The most progressively enriched category during lysosomal aging were proteins associated with the myelin sheath (**Figure 3h-i**). This included myelin components MOG, MOBP and PLP1 (**Figure 3d-f**, red). Myelin is rich in sphingolipids, and its increased endo/phagocytosis could contribute to the observed age-dependent increase in sphingolipids (**Figure 2**). Specific age-related changes also included altered abundance of many proteins involved in lysosomal storage diseases (LSDs, **Figure 3d-f**, bold). This included accumulation of prosaposin (PSAP) in the aged mice (18- and 24-months old). Mutations in the *PSAP* gene are observed in several LSDs, including Gaucher disease and metachromatic leukodystrophy, and are associated with increased predisposition to Parkinson’s disease. Other proteins accumulating in aged lysosomes included palmitoyl-protein thioesterase 1 (PPT1) and tripeptidyl peptidase 1 (TPP1), mutations in which genes are both associated with neuronal ceroid lipofuscinoses (CLN1 and CLN2, respectively)^49^.

### Data integration and identification of factors associated with lysosomal aging

We used factor analysis and machine learning (ML) approaches to integrate lysosomal lipidomic and proteomic data over time. After applying batch correction and outlier removal methods (**Supplementary Figure S4**), we used multi-omics factor analysis (MOFA) to capture shared variance across lipids and proteins over time. Changes in many proteins, lipids, or other biomolecules are correlated in high-throughput analyses. This correlation reflects alteration in underlying biological pathways that are not directly observable but give rise to the observed changes in individual protein and lipid species. By grouping shared variation over time into calculated factors, we were able to obtain a measurement of these biological pathways. MOFA resulted in ten factors that explained 78.6% of variance in the lipid data and 65.2% of variance in the protein data, with four factors explaining variance across both -omics types (**Figure 4a**).

**Figure 4.**
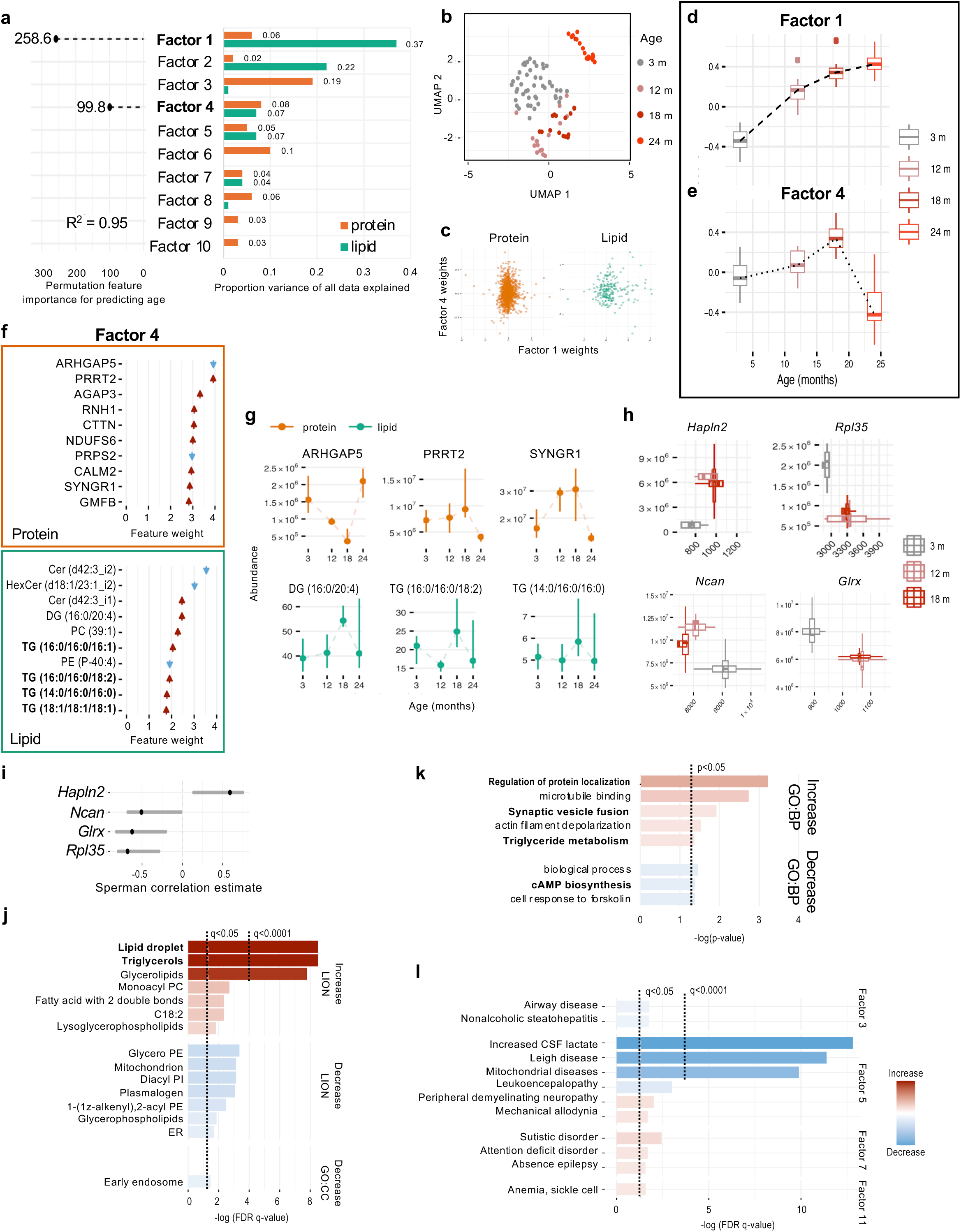
Multi-omics factor analysis (MOFA) of lysosomal aging. **a,** Multi-omics factor analysis of data results in ten factors that explain 78.6% of variance in the lipid data and 65.2% of variance in the protein data (right). factor 1 and factor 4 are chosen by ML feature selection as predictive of age. A random forest regression model R^2^ of 0.95 demonstrates that the model fitted on these two factors is a good predictor for sample age. **b**, Uniform manifold approximation (UMAP) embedding of ten multi-omics factors into two dimensions. Points generally cluster based on sample age. **c**, Plotting protein and lipid weights for factor 1 against those of factor 4 demonstrates that the factors have distinct identities. **d-e,** Factor scores for factor 1 and factor 4 plotted against age. Factor 1 increases monotonically with age (d); factor 4 increases up until 18-months followed by rapid decrease (e). **f**, Factor weights of the top 10 proteins and lipids associated with factor 4. A red arrow indicates increase with age, blue arrow indicates decrease with age. **g**, Abundance of selected individual proteins and lipids strongly associated with factor 4 show similar aging trends as the overall factor score. Vertical lines represent the interquartile range at each age point. **h**, Boxplots of protein abundance (ordinate) plotted against RNA expression (abscissa) for gene-protein pairs where abundance and expression are significantly correlated (*Hapln2*) or significantly anticorrelated (*Rpl35, Ncan, Glrx*). **i**, 95 % Confidence interval for Spearman correlation between expression and abundance values for each gene-protein pair. **j**, Enrichment-style analyses of factor 4 (vs all detected lipids or proteins) using LION lipid ontology and Gene Ontology. For clarity, Gene Ontology results have been filtered to the Cellular Component aspect and LION results have been filtered to child terms of “lipid classification”, “type by bond”, “cellular component”, and “fatty acid unsaturation”, excluding specific lipid chemical compositions. **k**, Gene Ontology Biological Process (GO:BP) analysis for factor 1 proteins. **l**, False discovery rate q-value from over-representation analysis plotted for five DisGenNet disease ontology terms for factors 3, 5, 7 and 11.

Projecting the ten factors onto two dimensions showed grouping based on age (**Figure 4b**). This was followed by application of supervised ML methods to determine which of the generated factors are relevant to age. Least absolute shrinkage and selection operator (lasso) regression demonstrated that two of the MOFA-identified factors (1 and 4) were best able to predict age in both lipids and proteins (combined *R*^2^ = 0.95 based on random forest regression model), with permutation feature importance stronger for factor 1 than for factor 4 (**Figure 4a**). Although there was some overlap between individual components, overall lack of strong correlation between factor 1 and factor 4 demonstrated that they are distinct (**Figure 4c**). Both factors increased with age, but with different kinetics: factor 1 captured near-monotonic effect increasing with age; factor 4 captured non-monotonic effect with increase up to 18 months, followed by sharp decline at 24 months (**Figure 4d-e**).

We applied the elbow method to determine a discrete set of proteins and lipids to examine in relation to factor 4, resulting in 256 proteins and 24 lipids (**Figure 4f, Supplementary Table S18**). We confirmed that changes in absolute abundance of both protein and lipid components over time were consistent with the overall factor 4 kinetics (**Figure 4g**). Performing correlation analyses between RNA-sequencing and protein abundance data, we found only a single gene, *Hapln2*, with mRNA pattern correlated with its corresponding protein and three genes, *Rpl35*, *Glrx*, and *Ncan*, demonstrating anticorrelated patterns (**Figures 4h-i**).

Applying lipid ontology (LION) to factor 4, we found the strongest positive association with neutral lipids including triacylglycerols (TAGs) and the lipid storage organelles, lipid droplets (**Figure 4j**). Accumulation of lipid droplets in microglia and neurons has been previously reported during brain aging and in neurodegenerative diseases and in microglia shown to be associated with age-related increase in neuroinflammation^50^. LION analysis of factor 4 also included a lesser association with increased lysophospholipid species and with linoleic acid (C18:2). Additionally, we found an association with decrease in overall glycerophospholipids and plasmalogens, as well as with mitochondria and endoplasmic reticulum associated lipids. Applying GO:CC enrichment analysis, we identified a single term, early endosome, significantly decreased with factor 4. Although Gene Ontology Biological Process (GO:BP) analysis of factor 4 proteins did not reach FDR significance threshold, *t*-test based enrichment revealed positive association with triglyceride metabolism, consistent with results of LION lipid analysis (**Figure 4k**). Transcription factor over-enrichment analysis for elbow method cutoff sets revealed no significant associations between factors 1 and 4 with any TFlink mouse transcription factors, although significant associations were found for factors 3 and 5.

With disease ontology over-representation analysis, we identified no significant associations between factor 4 and any diseases. We did, however, identify several disease associations with other factors (**Figure 4l**). Factor 5 was associated with 57 different disease ontology terms, although many were overlapping. Notable were decreases in Leigh disease (a rare inherited neurometabolic disorder), lactic acid, and mitochondrial disease-related proteins, and increases in peripheral demyelinating neuropathy and mechanical allodynia-related proteins. Factor 7 was associated with 3 different disease ontology terms, which were all increased: autistic disorder, attention deficit-hyperactivity disorder (ADHD), and absence epilepsy. Factor 10 was associated with an increase in proteins related to a single disease term, sickle cell anemia.

### Factor 1 analyses point to sphingolipid changes as drivers of lysosomal aging

MOFA identified factor 1 as the most significantly associated with age. Applying the elbow method cutoff to factor 1 resulted in associations with 101 proteins and 71 lipids (**Figure 5a, Supplementary Table S19**). Consistent with our pairwise analyses of proteomic data, factor 1 proteins with highest feature weights included age-dependent increase in myelin proteins such as MOG, enzymes associated with lysosomal storage diseases such as PPT1 and TPP1, and sphingolipid catabolism proteins such as ASAH1. Most important lipid features included lysosomal lipid BMP and multiple species of HexCer and sulfatides, all of which increased with age. Changes in the absolute abundance of both protein and lipid components over time were consistent with the overall factor 1 kinetics (**Figure 5b**). Performing correlation analyses between RNA-sequencing and protein abundance data, we found 12 out of 101 (11.9%) gene-protein pairs including *Ppt1*, *Asah1* and *Psap* (**Figure 5c**, **Supplementary Figure 5a-b**) with patterns significantly correlated between these two -omics types. Except for *Dpysl3*, *Ass1* and *Homer1* (2.97%), most of the time-dependent changes in gene expression were not in themselves significant (**Supplementary Figure S5c**). A single gene (*Rpl35*) demonstrated significant opposite pattern with the corresponding protein. Two additional gene-protein pairs including *Mog* also showed opposite pattern but did not reach correlation significance (**Figure 5c**, **Supplementary Figure S5d**). Performing over-enrichment analysis of gene-protein pairs with positive correlation, we found that 10 out of 12 play roles in cell projection or neuron projection, with the exceptions being *Psap* and *Aldh9a1* (**Supplementary Figure S5e**).

**Figure 5.**
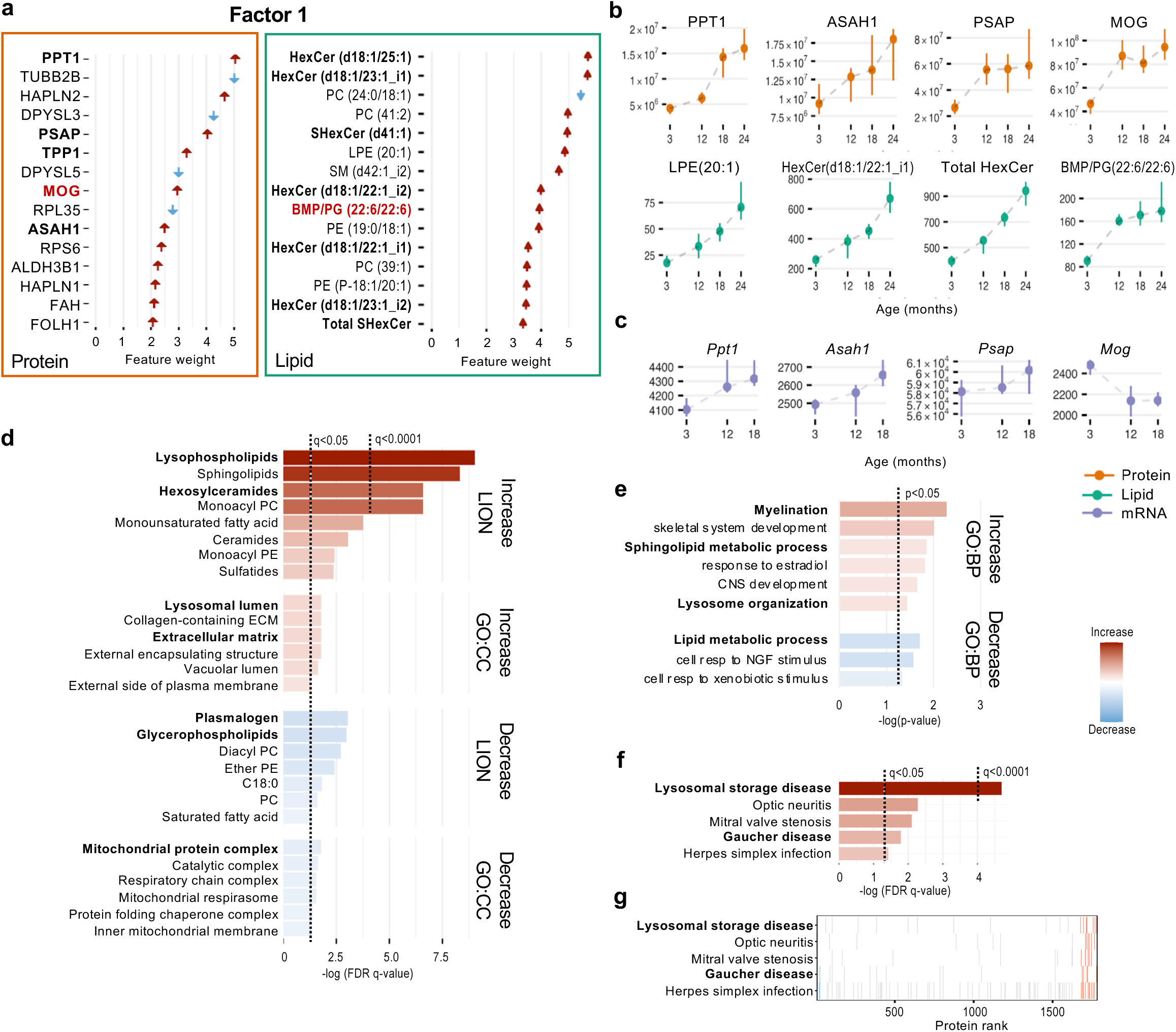
Analysis of lysosomal aging Factor 1 points to changes in sphingolipid metabolism. **a,** Factor weights of the top 15 proteins and lipids associated with factor 1. A red arrow indicates increase with age, blue arrow indicates decrease with age. **b**, Abundance of selected individual proteins and lipids strongly associated with factor 1 show similar aging trends as the overall factor score. Vertical lines represent the interquartile range at each age point. **c**, Trends in gene expression matched to proteins associated with factor 1 for three of the four time points. Some genes demonstrate the same trend with age as their protein counterparts, whereas others demonstrate the opposite trend. **d**, Enrichment-style analyses of factor 1 (vs all detected lipids or proteins) using LION lipid ontology and Gene Ontology. For clarity, Gene Ontology results have been filtered to the Cellular Component aspect and LION results have been filtered to child terms of “lipid classification”, “type by bond”, “cellular component”, and “fatty acid unsaturation”, excluding specific lipid chemical compositions. **e**, Gene Ontology Biological Process (GO:BP) analysis for factor 1 proteins. **f**, False discovery rate *q*-value from over-representation analysis plotted for five DisGenNet disease ontology terms. **g**, Barcode plots for corresponding proteins. Protein rank indicates the rank protein weight along factor 1. Solid lines indicate that a protein is annotated with a given disease ontology, with red lines corresponding to proteins passing the elbow method that are increased and blue lines corresponding proteins passing the elbow method that are decreased.

LION analysis of factor 1 revealed that it was dominated by increase in lysophospholipids, including both LPC and LPE species, and sphingolipids, including HexCer and sulfatides, but not sphingomyelins (**Figure 5d, Supplementary Figure S5f**). Similar to factor 4, levels of plasmalogens and overall phospholipids were decreased. Accumulation of lysophospholipids and concomitant decrease in phospholipids suggest age-related increase in phospholipid degradation by lysosomal phospholipases^40^. Additionally, cytoplasmic phospholipases such as cPLA2, can translocate to lysosomes and mediate hydrolysis of lysosomal membrane phospholipids leading to lysosomal dysfunction^16^. Overall decrease in plasmalogens has been previously noted during brain aging and in brain injury^42^. Plasmalogens are important components of cellular membranes where they regulate cellular signaling pathways and serve as antioxidants. ^43^ Their decrease may suggest increased sensitivity of aged lysosomal membranes to oxidative damage. Enriched LION terms also included increase in monounsaturated fatty acids and decrease in fully saturated fatty acid chains, suggesting increase in lysosomal membrane fluidity^51^.

The most prominent increased factor 1 components were sphingolipids, especially HexCer and sulfatides. Overall increase in sulfatide abundance was also one of the top individual factor 1 features associated with aging (**Figure 5a**). HexCer and sulfatides are highly abundant components of myelin, pointing to potential role of myelin as driver of the observed lysosomal lipid changes. This is consistent with previous reports indicating myelin phagocytosis as promoting microglial dysfunction during brain aging^18^. This was further supported by analysis of factor 1 proteins, which included increases in myelin components such as MOG, MBP, PLP1 and CLDN11. GO:CC analysis indicated overall decreases in terms related to intracellular compartments such as mitochondria and increase in terms related to extracellular localization and to lysosomal lumen (**Figure 5d**). This suggests increased delivery of extracellular as opposed to intracellular cargos to lysosomes in the aged brain and is consistent with previously reported age-dependent changes to autophagosomal cargo composition^39^. Although GO:BP analysis of factor 1 did not reach FDR significance threshold, *t*-test based enrichment pointed to importance of pathways related to myelination, sphingolipid metabolism and lysosome organization (**Figure 5e**).

We conducted disease ontology over-representation analysis, identifying five disease terms associated with factor 1. The most significant association was increase in lysosomal storage diseases (LSDs), followed by optic neuritis, mitral valve stenosis, Gaucher disease, and herpes simplex infections (**Figure 5f-g**). Many LSDs including Gaucher disease are caused by mutations in genes involved in lysosomal sphingolipid catabolism. Consistently, top-weighted factor 1 components included lipid metabolic enzymes and their co-factors (ASAH1, PPT1, TPP1 and PSAP), pointing to overall perturbation of lysosomal lipid metabolism in the aging brain.

### Lysosomal sphingolipid catabolism and myelin phagocytosis contribute to altered lysosomal lipidome

The overall major conclusions from factor 1 analysis were consistent with our pairwise lipid and protein comparisons and suggested that underlying causes of the observed age-dependent changes in lysosomal lipid and protein composition include 1) changes in lysosomal sphingolipid catabolism, and 2) increase in delivery of sphingolipid-rich myelin-derived phagocytic/endocytic cargo to the lysosomes^18^. To confirm these ML-based predictions, we performed *in vivo* validation studies.

BMP is a lysosomal lipid involved in regulation of lipid catabolism by serving as anchor and cofactor for several degradative enzymes^41^. We used imaging to confirm age-dependent accumulation of BMP and demonstrate that both neuronal and microglial cells were affected (**Figure 6a-c, Supplementary Figure S6a-b**). To make sure accumulation of BMP was occurring specifically in lysosomes, we used super-resolution stimulated emission depletion (STED) imaging, which demonstrated that BMP signal in the aged brains was contained within lysosomes (**Figure 6d-e**).

**Figure 6.**
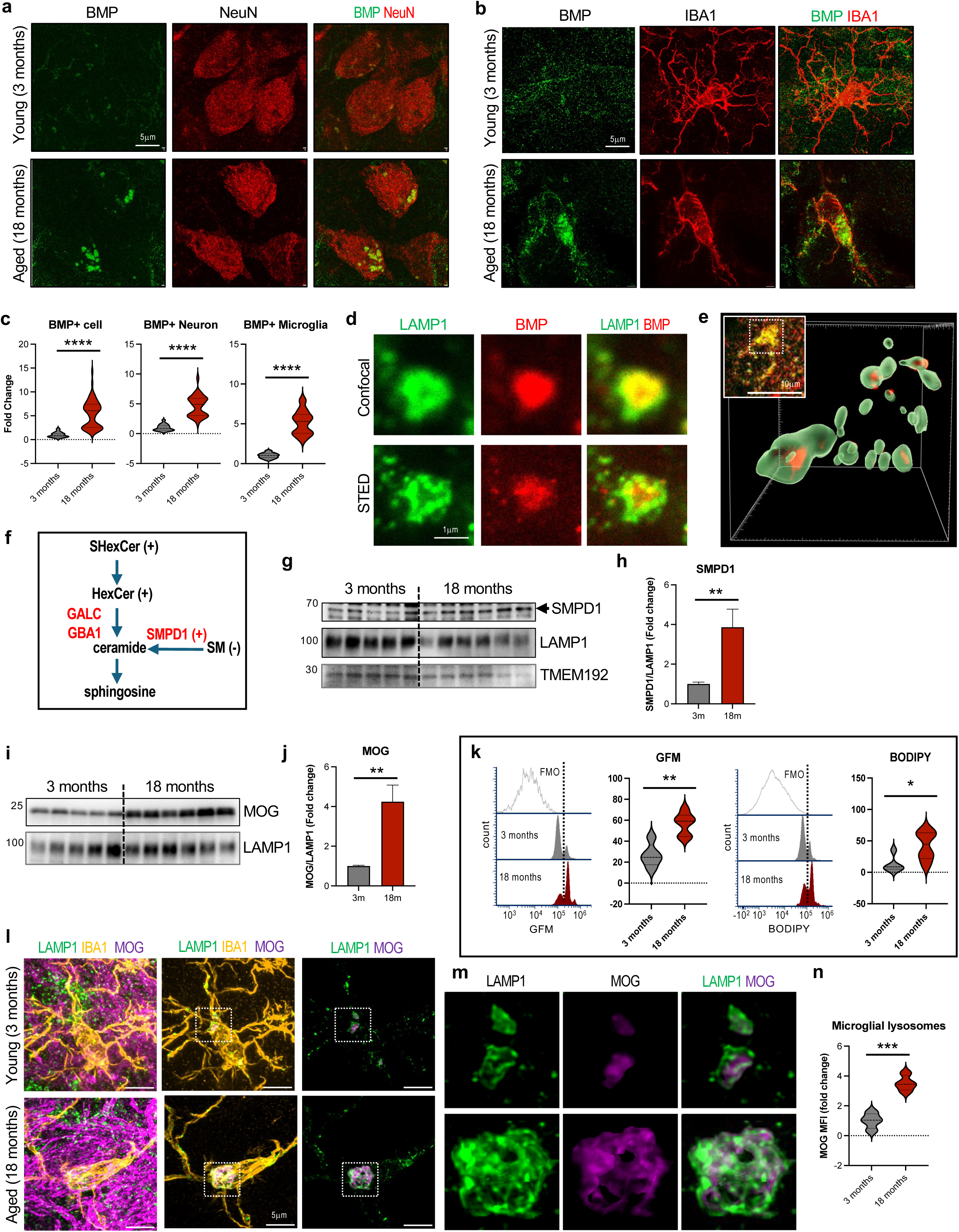
Lysosomal sphingolipid metabolism is altered during brain aging. **a-b,** Airyscan immunofluorescence images demonstrating age-dependent accumulation of BMP in neurons (a) and microglia (b). **c**, Quantification of data from a and b. \*\*\*\**p*-value < 0.0001, unpaired t-test with Welch’s correction; n = 4-5. **d-e**, STED immunofluorescence images demonstrating that BMP is localized within lysosomes. Comparison between confocal and STED images (d) 3-D rendering of neuronal lysosomes containing BMP (e) are shown. **f**, Schematic summary of lysosomal sphingolipid catabolism pathway. Sphingolipid species are in black, catabolic enzymes in red. +/- indicate direction of age-dependent change detected in lipidomic/proteomic analyses. **g-h**, Western blots and quantification of age-dependent changes in sphingolipid catabolic enzymes in cortical lysosomes. Data are mean +/- SEM; *p-value < 0.05, **p-value < 0.01; Mann Whitney test; n = 6. **i-j,** Western blots and quantification of age-dependent changes in myelin protein MOG in cortical lysosomes. **k,** Flow cytometry analyses and quantification demonstrating higher accumulation of myelin (green FluoroMyelin, GFM) and neutral lipids (BODIPY) in microglia from aged as compared to young cortex. **l-n,** Immunofluorescence images demonstrating age-dependent accumulation of MOG in microglial lysosomes. Middle and right column images depict LAMP1 and MOG signal restricted to IBA1^+^ mask for visualization and quantification. Closeups of indicated areas are shown in m. ***p-value < 0.001, unpaired t-test; n = 4

Our MOFA-downstream analyses indicated that changes in abundance of lysosomal enzymes involved in lipid catabolism could be involved in the observed perturbation of sphingolipid levels (**Figure 6f**). One of the proteins identified as increased with age was sphingomyelin phosphodiesterase SMPD1, an enzyme which degrades lysosomal sphingomyelin to ceramide. SMPD1 interacts with BMP^52^ and it activation could lead to the observed decrease in sphingomyelin levels, despite concurrent increase in other sphingolipid species. We confirmed increased SMPD1 protein abundance in lysosomes purified from the aged as compared to young brains (**Fig 6g-h**). Total tissue levels of SMPD1 protein were also increased with age, suggesting regulation at both overall protein abundance and subcellular localization levels (**Supplementary Figure S6c-d**). Lysosomal HexCer are degraded to ceramides by galactosylceramidase (GALC) and glucosylceramidase (GBA1, **Figure 6f**)^53,54^. Consistent with the possibility that HexCer and sphingomyelin degradation may be differentially regulated during brain aging, we did not observe significant changes in lysosomal abundance of either enzyme in either proteomic or western blot analyses, although, tissue abundance of GALC 50 kDa subunit was increased at 18 months (**Supplementary Figure S6e-h**). Activities of sphingolipid metabolism enzymes are regulated by interaction with other proteins including cleaved fragments of the PSAP protein, saposins A through D. While our proteomic data identified PSAP as one of the lysosomal proteins most increased with age, western blot analyses failed to detect changes in full length PSAP (**Supplementary Figure S6c-d**). To account for this discrepancy, we re-analyzed our proteomic data and found that only saposins A and D increased with age, and that saposin D, which has been reported to preferentially activate SMPD1, was the most abundant (**Supplementary Figure S6i**)^55^. Saposin C, the preferential cofactor for GBA1, was not changed^56^. Gene expression corresponding to any of the sphingolipid catabolic enzymes was not altered (**Supplementary Figure S6j**).

Factor 1 analyses also pointed to myelin as the likely source of sphingolipids accumulating in the aged lysosomes. We confirmed increased abundance of the myelin protein MOG in the aged as compared to young lysosomes and tissues (**Figure 6i-j**, **Supplementary Figure S6k-l**). We hypothesized that increased myelin phagocytosis by microglia could lead to the observed increase in lysosomal sphingolipids^18^. Consistently, our flow cytometry analyses indicated that more CD11B^+^ microglia from the aged brains were positive for FluoroMyelin (**Figure 6k**, **Supplementary Figure S6m**). Additionally, we observed more microglial neutral lipid accumulation (BODIPY) at 18 months (**Figure 6k**), consistent with factor 4 prediction and previous reports of microglial lipid droplet accumulation in the aged brain ^50^. We used imaging to confirm MOG accumulation in aged IBA1^+^ microglia (**Figure 6l**) and demonstrate that it specifically localized to the lysosomes (**Figure 6m-n**, **Supplementary Figure S7a-c**).

### Age-related lysosomal changes lead to lysosomal and autophagy dysfunction

Accumulation of sphingolipids including HexCer has been shown to cause lysosomal dysfunction in lysosomal storage diseases^27,57^. We used super resolution Airyscan imaging to compare lysosomal compartment size and morphology in young versus aged cortex. Our data demonstrated overall increase in lysosomal volume and in abundance of very large (above 0.5 μm^3^) lysosomes in both neurons and microglia in the aged as compared to young cortex (**Figure 7a-f, Supplementary Figure S8a-b**). These changes were particularly pronounced in microglia, consistent with age-related accumulation of myelin detected by flow cytometry (**Figure 6k**) and previously reported effect of myelin on microglial aging^13,18^.

**Figure 7.**
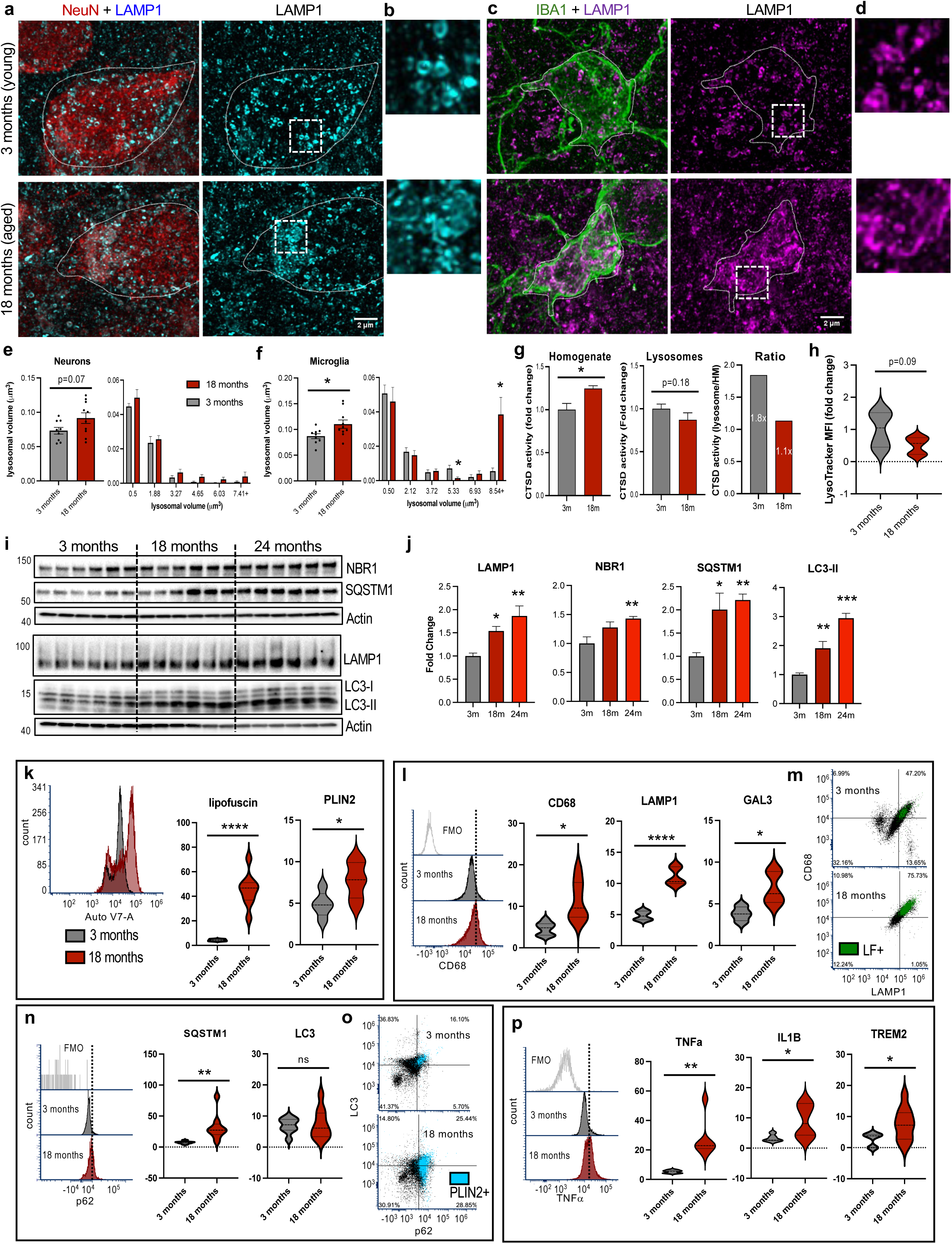
Lysosomal lipid changes lead to lysosomal dysfunction during brain aging. **a-e,** Airyscan immunofluorescence images demonstrating increased lysosomal size in neurons (a-b) and microglia (c-d) of aged as compared to young mouse cortex. Images in b and d are closeups of indicated areas in a and c, respectively. **e-f,** Quantification of total lysosomal volume and lysosomal size distribution in neurons (e) and microglia (f). **g,** Quantification of CTSD enzymatic activity in cortical homogenates (HM) and lysosomal fractions from young vs. aged mice. **h,** Quantification of LysoTracker-Red intensity in lysosomal fractions. i-j, Western blot and quantification of autophagy-lysosomal pathway proteins p62, NBR1, LC3 and LAMP1. **k-o,** Flow cytometry analyses and quantification demonstrating accumulation of lipid (lipofuscin and lipid droplets marked with PLIN2) in microglia from aged as compared to young cortex (k) and their correlation with lysosomal expansion (CD68, LAMP1) and damage (GAL3, l-m) and with inhibition of autophagy (SQSTM1, LC3, n-o). **p,** Flow cytometry data demonstrating increased microglial inflammation in the aged brain. Bar graphs are mean +/- SEM; *p-value < 0.05, **p-value < 0.01, ***p-value < 0.001, ****p-value < 0.0001; unpaired t-test (e-h, k-p), multiple comparison t-test (size distribution in e-f) or one-way ANOVA (j); n=5-6

Lysosomal enlargement and accumulation of undigested cargo are often observed during lysosomal dysfunction, suggesting that the aged lysosomes may be less active^16^. Consistent with overall increased abundance of lysosomal proteins in the aged brain, we observed higher activity of lysosomal enzyme CTSD in brain homogenates from 18-as compared to 3-month-old mice. Conversely, CTSD activity was lower in purified lysosomes from aged versus young mice. Although this did not reach significance, the ratio of lysosomal to tissue CTSD activity was considerably lower in aged as compared to young mice (**Figure 7g**). Additionally, lysosomes from aged mice showed poor acidification, as assessed by decreased LysoTracker intensity (**Figure 7h**, **Supplementary Figure S8c**).

To obtain a functional measure of lysosomal activity we looked at the levels of autophagy, a lysosome-dependent catabolic pathway known to be inhibited in many neurodegenerative diseases and reported to be down-regulated during brain aging^20,24^. Consistent with age-related decline in lysosomal function, we observed signs of progressive autophagy inhibition including accumulation of autophagy cargo receptors NBR1 and SQSTM1 and increase in autophagosome-associated LC3-II (**Figure 7i-j**). None of the changes were detected at the transcriptional level (**Supplementary Figure S9a**). Positive autophagy regulators BECN1 and ULK1 were increased with age, suggesting that inhibition of autophagy was not due to upstream inactivation of its regulatory pathways (**Supplementary Figure S9b-c**). We also did not observe decrease in activity of the mTORC1 kinase, a negative upstream regulator of autophagy.

Lipid accumulation has been implicated in microglial dysfunction and reprogramming in the aging brain. This included presence of lipofuscin, a lipid byproduct, in the endo-lysosomal compartments and microglial accumulation of lipid droplets^18,50^. Our flow cytometry analyses confirmed accumulation of both lipofuscin and lipid droplets (PLIN2) in aged mouse microglia (18 months) as compared to young controls (**Figure 7k**)^58^. Microglial lipid changes were accompanied by increase in lysosomal markers LAMP1 and CD68 consistent with lysosomal expansion observed by IF, and accumulation of the marker of lysosomal damage, GAL3 (**Figure 7l**). Consistent with the proposed role of lipid accumulation in lysosomal dysfunction, lysosomal changes were preferentially observed in microglial cells accumulating high levels of lipid (**Figure 7m**). Changes in lysosomal markers were not explained by altered gene expression (**Supplementary Figure S9a, d**). We also observed age-dependent inhibition of microglial autophagy as indicated by accumulation of SQSTM1 (**Figure 7n**), which was preferentially detected in lipid-accumulating cells (**Figure 7o**). We previously demonstrated that inhibition of microglial autophagy leads to increased inflammation^59^. Consistently, microglia from the aged brains were more proinflammatory (**Figure 7p, Supplementary Figure S9e**).

## Discussion

Recent data indicate that changes in lipid composition and distribution affect brain aging and predisposition to neurodegenerative diseases. However, the identity of the affected lipids and the functional *in vivo* consequences of these alterations remain poorly understood. Here we evaluated the role of lysosomal lipids during brain aging by analyzing lipid and protein composition of lysosomes isolated from the mouse cortex from 3- to 24-months of age and using ML approaches to integrate and interpret these longitudinal data. This was followed by experimental *in vivo* validation of the ML predictions. Our analyses indicate that mechanisms underlying lysosomal aging include combination of age-dependent increase in lysosomal delivery of sphingolipid-rich myelin components and changes in lysosomal sphingolipid catabolism favoring degradation of sphingomyelin over HexCer. The overall result is pronounced accumulation of HexCer, which resembles what is observed in lysosomal storage diseases (LSDs), and leads to lysosomal dysfunction and inhibition of lysosome-dependent cellular processes including autophagy.

Although technological advances allow generation of individual data sets to describe large-scale changes in biomolecule abundance (e.g., transcriptome, proteome, lipidome), integration of the various “omics” data has remained challenging^60–62^. This presents a potential limitation, as understanding of many pathways requires consideration of interactions between different types of molecules, such as influence of catabolic enzymes on lipid abundance and the role of lipids in regulation of protein function^63^. Compared to genomics and transcriptomics, integration of proteomic and lipidomic data poses additional unique challenges^62^. The proteome bridges the genotype and gene expression to the phenotype of the cell and organism. However, several studies including data presented here, demonstrate poor agreement between transcriptomic and proteomic data, indicating contribution of posttranscriptional factors^44,45^. Lipidome is not directly encoded in the genome but rather is the result of interactions between protein function and the environment^62^. As a result, while the proteome and lipidome reflect phenotype more closely than the genome and transcriptome, their study is complicated by the lack of linear relationships. Furthermore, unlike DNA and mRNA, for proteins and lipids intracellular localization must be considered when establishing functional networks^16,62^. Many lipid and protein species dynamically reside in multiple locations, and their cellular distribution is important for specific interactions and biological function^15^. To overcome these challenges, we used a combination of biochemical organelle purification, integrated longitudinal MS-based multi-omics, factor analysis and application of ML to generate testable hypotheses regarding mechanisms contributing to lysosomal aging. This was followed by *in vivo* validation of the ML predictions. The resulting organelle-specific analytical-computational-experimental validation pipeline has allowed us to identify novel mechanisms of lysosomal dysfunction in the aging brain. We expect this approach can be applied to understanding mechanisms of organelle-specific changes in other diseases and models. Further refinements could be gained from combining organelle- and cell-type specificity, for example using transgenic mice expressing tagged organelles in a cell-type specific manner^64,65^.

Our analyses identified two factors encompassing both protein and lipid components and able to predict sample age. Factor 1, which increased linearly over time and was the best predictor of age, indicated lysosomal accumulation of myelin components including sphingolipids as underlying lysosomal aging. Factor 4 was non-monotonous and increased until 18 months of age, followed by a steep decline. Prominent lipid components of factor 4 were triglycerides. Accumulation of triglycerides and other neutral lipids has been previously shown to occur in the aging and neurodegenerative disease microglia and contribute to their dysfunction^50,66^. However, these reports described accumulation of lipids in the cellular lipid storage organelles, the lipid droplets (LDs)^50^. Our data indicate that in addition to LDs, triglycerides also accumulate in the aged lysosomes. Neutral lipids including cholesteryl esters and triglycerides are hydrolyzed by the lysosomal acid lipase (LAL), which deficiency can lead to lysosomal lipid accumulation in Wolman disease^67^. Our data suggest that decrease in LAL activity may also be a part of age-related lysosomal dysfunction, although further experiments will be needed to test this. The proximal causes of factor 4 kinetics will also require further investigation. The factor 4 peak at 18 months coincided with the onset of deficits in brain function, based on decline in memory and motor performance. We speculate that peak of factor 4 may reflect a switch from compensatory mechanisms supporting continued brain function during early stages of aging to overt decline later on.

Factor 1 pointed to myelin as the source of sphingolipids accumulating in the aged lysosomes. Myelin phagocytosis has been previously implicated in accumulation of lipofuscin, a lipid byproduct, in the endo-lysosomal compartments of aging microglia^18^. Additionally, increased levels of myelin in the white matter have been suggested to promote formation of a unique microglial population, the white matter associated microglia (WAM) in the aged and AD model mouse brain^13^. Our data further support the idea that myelin phagocytosis is one of the driving forces underlying age-related microglial dysfunction, and that this is mediated, at least in part, by myelin-derived lysosomal lipid accumulation. Additionally, we identified the specific lipids and lipid classes involved, most prominently HexCer and sulfatides.

Sphingolipids, including HexCer, sulfatides and sphingomyelins are all prominent components of myelin. However, unlike HexCer, we did not observe accumulation of sphingomyelin in the aged lysosomes. Our analyses indicate that brain aging is associated with altered lysosomal sphingolipid metabolism favoring degradation of sphingomyelin over HexCer. This may be in part the result of increased abundance of sphingomyelin degrading enzyme SMPD1 in the aged lysosomes. Additionally, lysosomal enzymes are activated by both lipid and protein cofactors. We identified BMP, a lipid anchor and cofactor for several degradative enzymes including SMPD1^41^, as factor 1 component significantly increased with age. BMP is negatively charged in the acidic pH of the lysosomes, which promotes its interaction with positively charged catabolic enzymes such as SMPD1^68^. SMPD1 binding to BMP within the lysosomes is required for sphingomyelin membrane extraction and degradation^52,69^. Although it is not clear why levels of BMP are increased with age, in a mouse model of macular degeneration, accumulation of lipofuscin species has been shown to trap BMP in the lysosomes, leading to SMPD1 activation^70^. Since our and published data demonstrate accumulation of endo-lysosomal lipofuscin in the aged mouse brain, we expect that similar mechanisms may play a role.

Another identified factor 1 component was PSAP. Polymorphisms in the *PSAP* gene are observed in both lysosomal storge and neurodegenerative diseases including frontotemporal dementia and PD^71^. PSAP protein is cleaved into several fragments (saposins A-D) which interact with different lysosomal catabolic enzymes to regulate their activity. Our proteomic data indicate that the only PSAP fragments increased with age are saposins A and D, with saposin D the most abundant. Although saposins show some promiscuity in their interactions, saposin D has been reported to preferentially activate SMPD1^55^. Conversely, saposin C, which is required for activation of GBA1 and degradation of glucosylceramides^56^, was not changed. Together these data indicate that increased abundance and activation of SMPD1 promote degradation of sphingomyelins but not HexCer during brain aging, leading to specific HexCer accumulation and overall lysosomal sphingolipid imbalance.

Reduction in autophagy has been observed during aging in diverse organisms from worms to flies to mammals, including humans^72^. Although expression of some autophagy genes has been reported to decline with age^24^, the full reasons for the age-dependent decrease in autophagy levels are not clear. Accumulation of HexCer has been linked to lysosomal dysfunction and inhibition of autophagy in lysosomal storage diseases^27,73,74^. Many forms of these diseases include CNS manifestations such as neurodegeneration, dementia and/or motor dysfunction. Polymorphisms in some of the LSD genes are also associated with predisposition to various neurodegenerative diseases, such as Parkinson’s disease (*GBA1* and *GRN*), Lewy body dementia (*GBA1*) and frontotemporal dementia (*GRN* and *PSAP*)^54,73,75,76^. Our data demonstrating accumulation of HexCer and sulfatides in the aged brain lysosomes suggest that similar mechanisms may contribute to inhibition of autophagy-lysosomal function during normal brain aging. The link between brain aging and lysosomal storage diseases is further supported by the age-related changes in lysosomal abundance of proteins implicated in LSDs, including PSAP, PPT1 and TPP1.

Because of its high lipid content and unique composition, changes in lipid metabolism have profound influence on the brain function, aging and predisposition to disease. However, connecting alterations in brain lipidome to it effect on cellular, tissue and organismal function remains challenging. The data presented here underscore the significance of organelle-specific analyses and longitudinal integration of multi-omics for elucidating lipid-dependent mechanisms pertinent to brain aging. Our data indicate that factors contributing to lysosomal aging resemble those observed in lysosomal storage diseases, adding a mechanistic link to the previously noted genetic connection between LSDs and neurodegenerative diseases.

## Author contributions

Project conceptualization and funding: MML, MPC, MAK and JWJ; experimental design: MML, MPC, MAK, JWJ, MK, TAB, CS and DPHN; experimentation: CS (animals and biochemistry), DPHN (imaging), MMW (proteomics), YM (lipidomics), AMT (flow cytometry), NG (biochemistry and imaging), OPR (behavior), NH (flow cytometry and behavior), ST (flow cytometry), SAK (lipidomics), SB (behavior), SZK (proteomics), CW (proteomics), NL (biochemistry) and MK (imaging); data analysis: CS (biochemistry), YC (statistics and machine learning), DPHN (imaging), MMW (proteomics), RTC (machine learning), SDK (imaging), CM (gene expression), JWJ (lipidomics) and MML (flow cytometry and behavior). MML drafted the manuscript with input from all the authors.

## Acknowledgements

This project was supported by R01NS115876 to MML, R33AG076858 to MML, MAK and MPC, R56AG081262 to MML, University of Maryland System AIM-HI Challenge Award to MML, MAK, JWJ and MPC. Confocal and STED imaging was performed at the University of Maryland Baltimore (UMB) Center for Innovative Biomedical Resources (CIBR) Confocal Microscopy Core Facility, supported by S10OD026698 and UM-MIND. Flow cytometry was performed at the CIBR Flow Cytometry Shared Service of the UMB Marlene and Stewart Greenebaum Comprehensive Cancer Center, supported by funds through the Maryland Department of Health’s Cigarette Restitution Fund Program (CH-649-CRF) and the NCI Cancer Center Support Grant (P30CA134274). RNA sequencing was performed by Maryland Genomics at the UMB Institute for Genome Sciences. The authors thank Dr. Xiaoxuan Fan for help with flow cytometry panel design and Dr. Luke Talon for advice on experimental design for RNA-seq.

---

**Figure S1.**
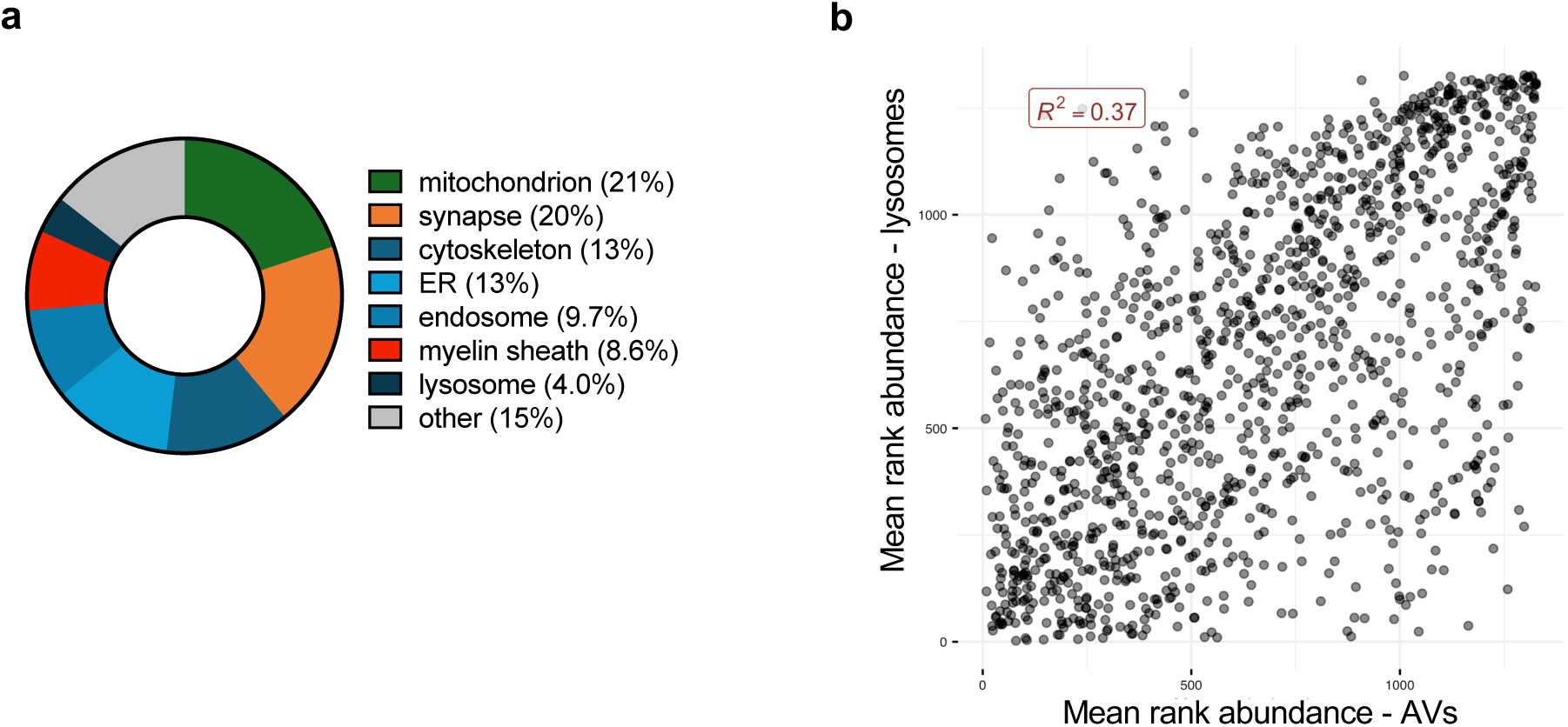
Proteomic characterization of mouse brain lysosomes. **a,** Summary of percent composition of proteins detected in purified lysosomal fractions from the mouse brain based on Gene Ontology Cellular Compartment (GO:CC) analysis. **b**, Mean rank abundance of each protein species in adult (3-month-old) mouse lysosomes plotted against that in adult (6-month-old) mouse autophagosomes (AV). Square of the Pearson correlation is reported in the top left (Kallergi et al., 2023, Neuron 111:2329–47).

**Figure S2.**
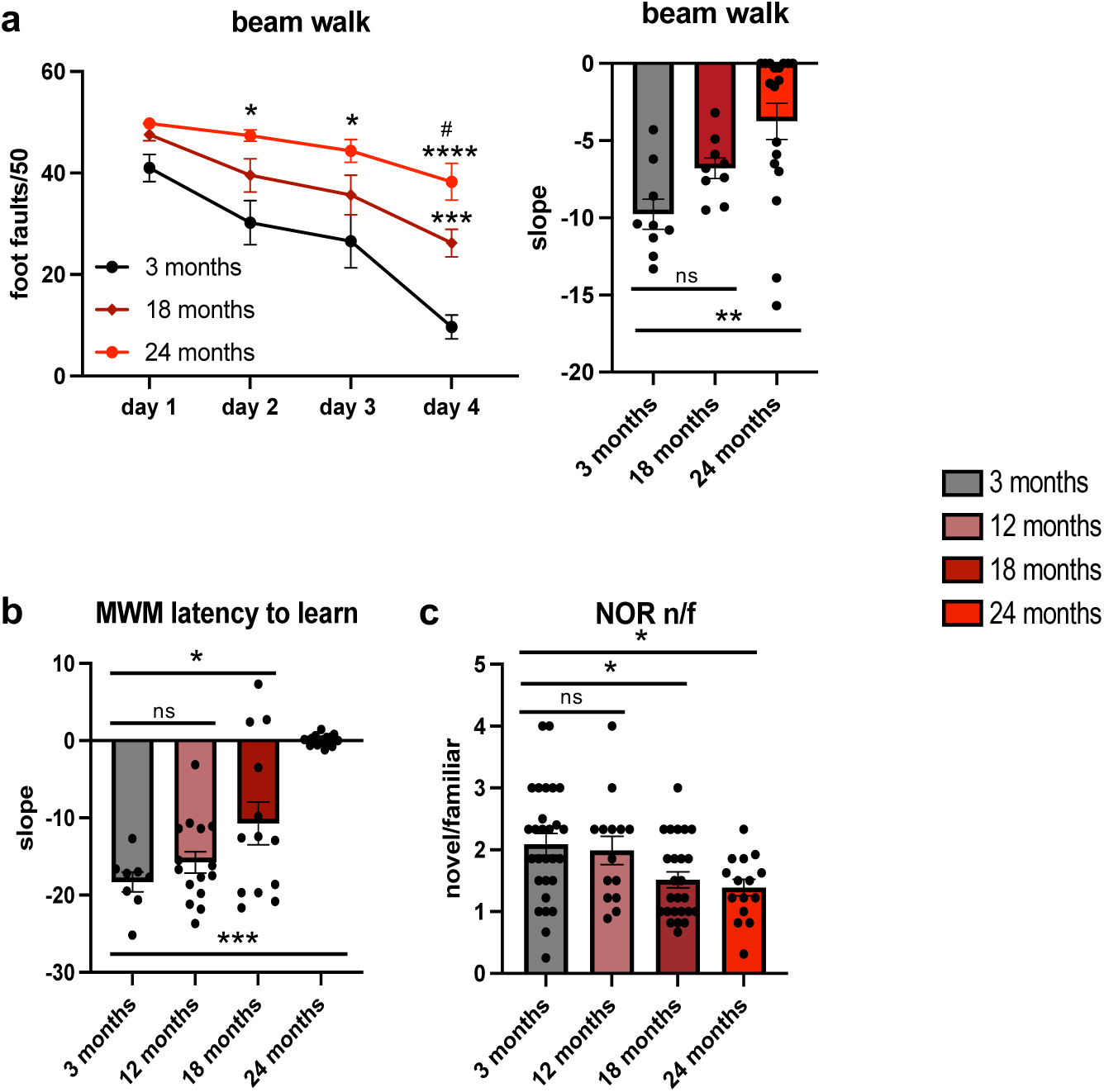
Progressive functional decline in the aging mice. **a,** Assessment of motor function by beam walk. Left: number of foot faults on training days 1-4 in different age groups. ^#^*p*-value < 0.05 for 18-vs 24-month-old; two-way RM ANOVA + Sidak’s multiple comparison test. Right – comparison of slope of the motor learning curve (decrease in foot faults) between age groups. n = 9-18. **b,** Assessment of spatial memory by Morris Water Maze. Comparison of the slope of the learning curve (over 4 days) is shown. n = 8-18. **c,** Assessment of non-spatial declarative memory by novel object recognition. Ratio of the time spent with novel versus familiar object (24 hours after familiarization) is shown. n = 14-28. All data are mean +/- SEM; dots correspond to scores for individual mice. *p<0.05, **p<0.01, ***p<0.001, ****p<0.0001 as compared to 3-month-old; one-way ANOVA + Tukey’s multiple comparison test.

**Figure S3.**
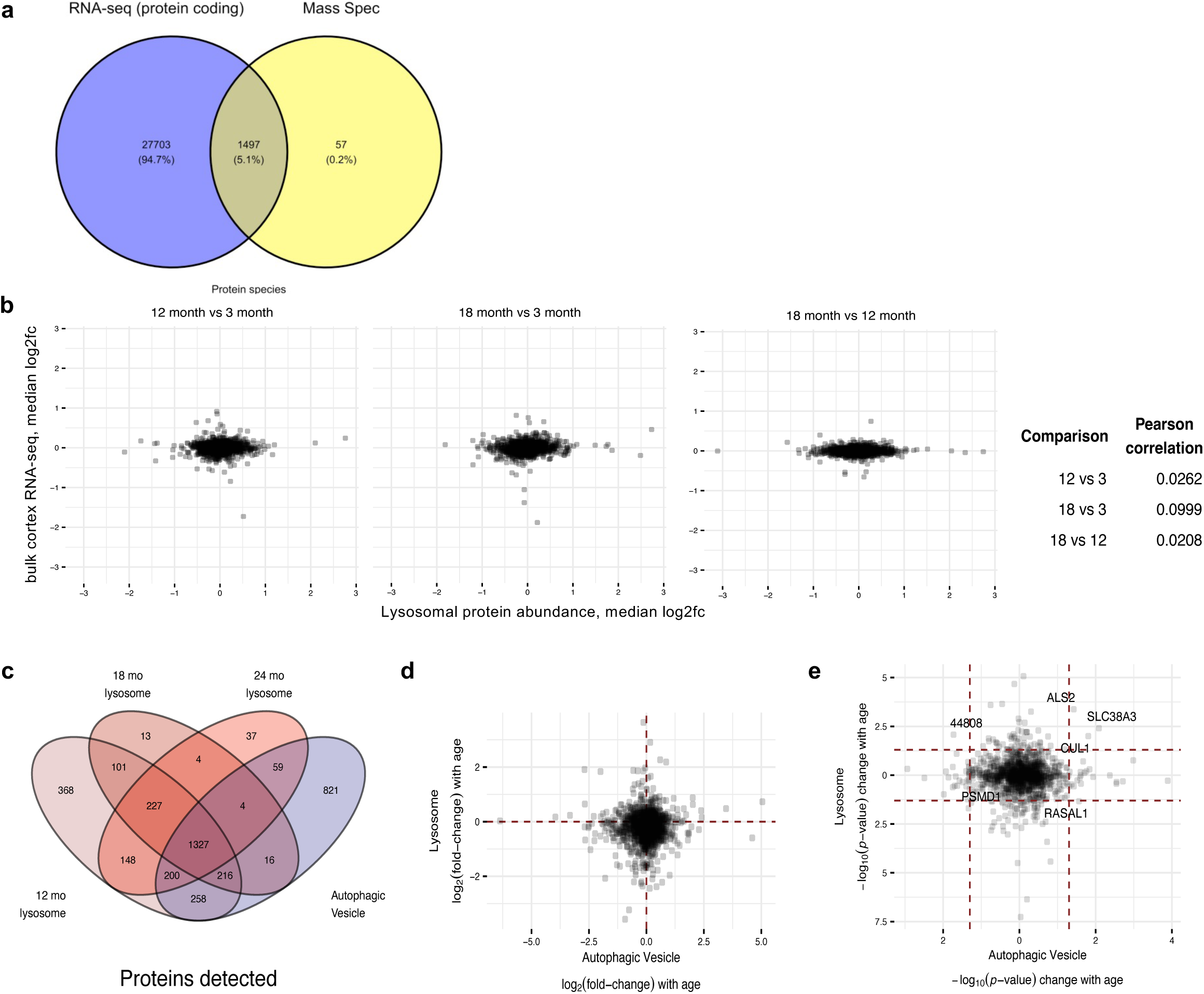
Comparison of lysosomal protein abundance to gene expression and autophagosomal protein abundance in the aging brain. **a,** Of the 1554 protein species detected by MS-based lysosomal proteomics, almost all (except 57) have corresponding genes represented in the bulk cortical RNA-sequencing data. **b,** Median log_2_ fold-change between different age points for RNA-sequencing data plotted against corresponding median log_2_ fold-change in protein mass spectrometry data. Each point represents a different protein/gene species. All proteins detected in both protein mass spectrometry and RNA-seq are plotted. Table of Pearson correlation values for each pairwise age comparison is included on the right. The lack of strong correlation between lysosomal protein data and cortical gene expression data demonstrates the difference in biological phenomena captured by the different analysis methods and the sampling between different compartments. **c,** Comparison of number of protein species detected in three batches of lysosome data by mass spectrometry with number of protein species detected in autophagic vesicles (Kallergi, et al., *Neuron*, 2024). **d**-**e**, Change in protein abundance between aged vs young mouse brain autophagic vesicles (abscissa) and aged vs young mouse brain lysosomes (ordinate). For autophagic vesicles, 6-month-old mice are used for the young time point and for lysosomes, 3-month-old mice are used. In both data sets, aged mice refer to 18-month-old samples. Points to the right of the *y*-axis or above the *x*-axis represent proteins that are increased in aged mice compared to young mice. In **d**, log_2_ fold-change with age is plotted, and in **e**, log_10_ *p*-value is plotted. *p*-values are calculated using a two-sided Student *t*-test. Six proteins with nominal *p*-values < 0.05 in both data sets are labeled with protein name or Entrez id.

**Figure S4.**
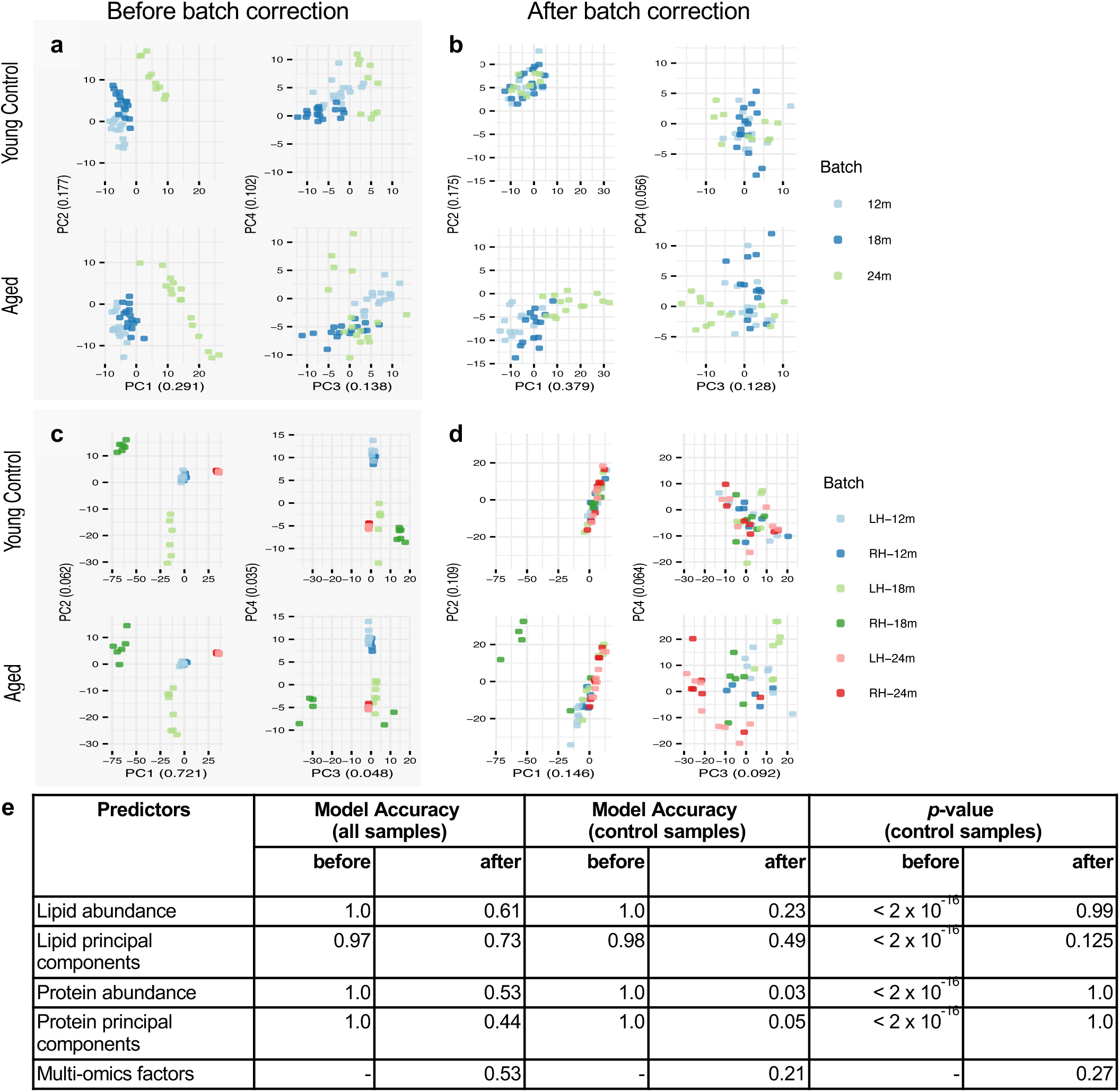
Lipid and protein data batch effect assessment using principal component analysis. **a-b,** Lipid batch effects. Top row of plots shows results for young control (3-month-old) mice. Bottom row of plots show results for aged mice. **a**, First four principal components (PCs) plotted prior to batch correction. PC1 and PC3 appear to separate 24-month batch control mice from 12- and 18-month batch. PC2 appears to separate 12-, 18-, and 24-month batches on a gradient. **b**, First four principal components plotted after batch correction. None of the PCs plotted visually separates the control mice by batch. PC1 appears to correspond to age in aged mice, whereas PC2 separates young mice from aged mice. **c-d**, Protein batch effects. Top row of plots shows results for young control (3-month-old) mice. Bottom row of plots show results for aged mice. LH corresponds to left-hemisphere batch/samples and RH corresponds to right-hemisphere batch/samples. **c**, First four principal components plotted prior to batch correction. All four PCs appear to distinguish young control samples from different batches in some way. **d**, First four principal components plotted after batch correction. None of the PCs plotted visually separates the control mice by batch. PC1 and PC2 appear to primarily distinguish four 18-month right hemisphere aged samples from all other aged samples. PC3 and PC4 appear to distinguish the aged samples by age. **e,** Machine learning model performance for predicting batch labels using various predictors before and after batch correction. Batch labels are modeled using classification random forest and *p*-values are reported comparing model accuracy on young control samples to the no information rate.

**Figure S5.**
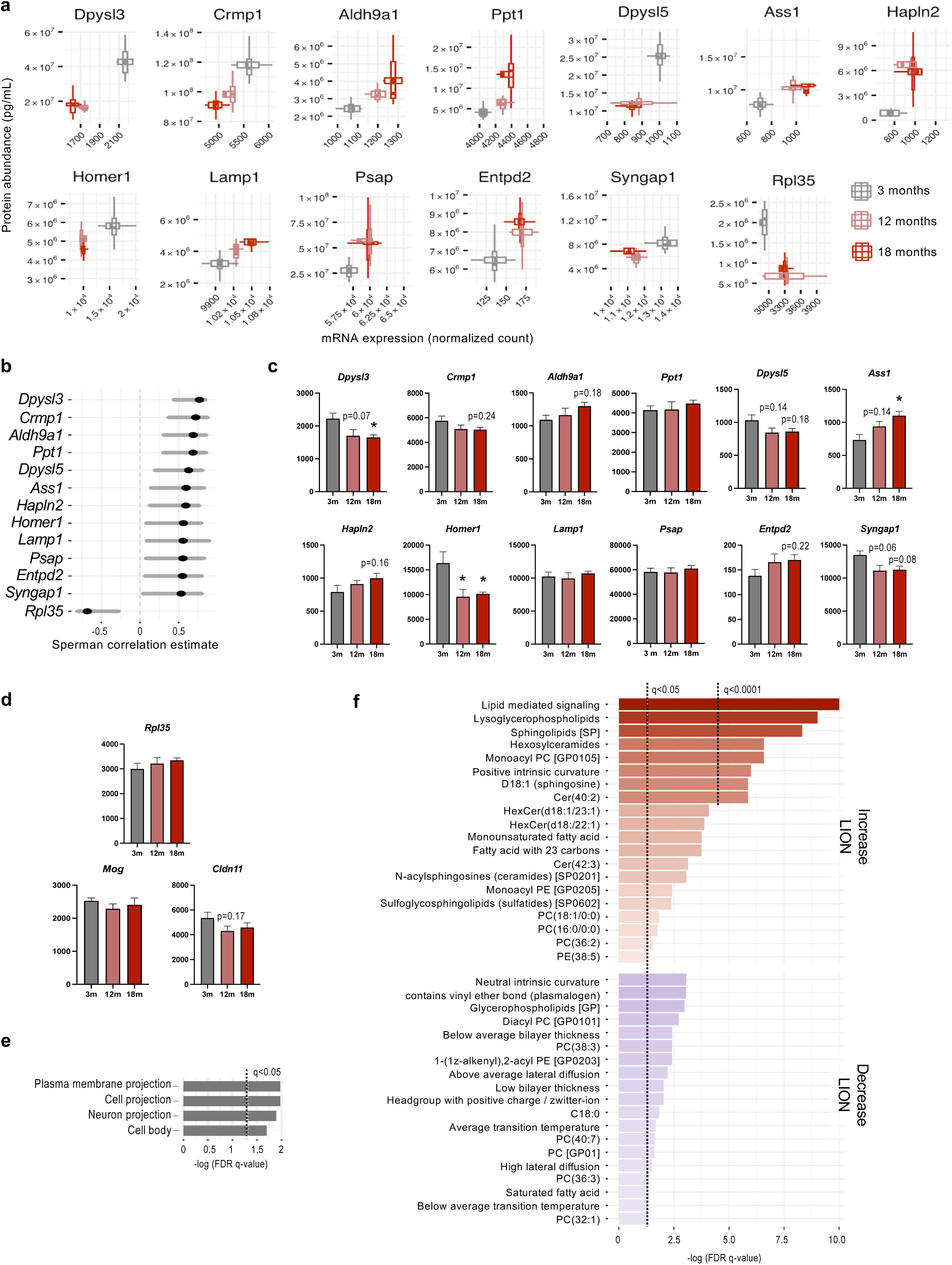
Additional analyses of factor 1 for protein abundance, RNA expression, and lipid ontology. **a,** Boxplots of protein abundance (ordinate) plotted against RNA expression (abscissa) for gene-protein pairs where abundance and expression are significantly correlated (*Dpysl3* through *Syngap1*) or significantly anticorrelated (*Rpl35*). **b**, 95 % Confidence interval for Spearman correlation between expression and abundance values for each gene-protein pair. **c-d**, RNA-seq based analysis of age-related changes in gene expression in cortical lysates between 3 and 18 months. Plots for genes significantly correlated with protein abundance are shown in (c), genes anti-correlated with protein abundance are in (d). Correlation for *Mog* and *Cldn11* did not reach significance. \**p*-value < 0.05; 1-way ANOVA + Dunnett’s multiple comparison test; n = 4. All *p*-values less than 0.25 are shown. **e**, Over-representation analysis of 12 expression-abundance correlated gene-protein pairs against background of all proteins associated with factor 1. **f**, Full LION lipid ontology analysis for factor 1 lipids.

**Figure S6.**
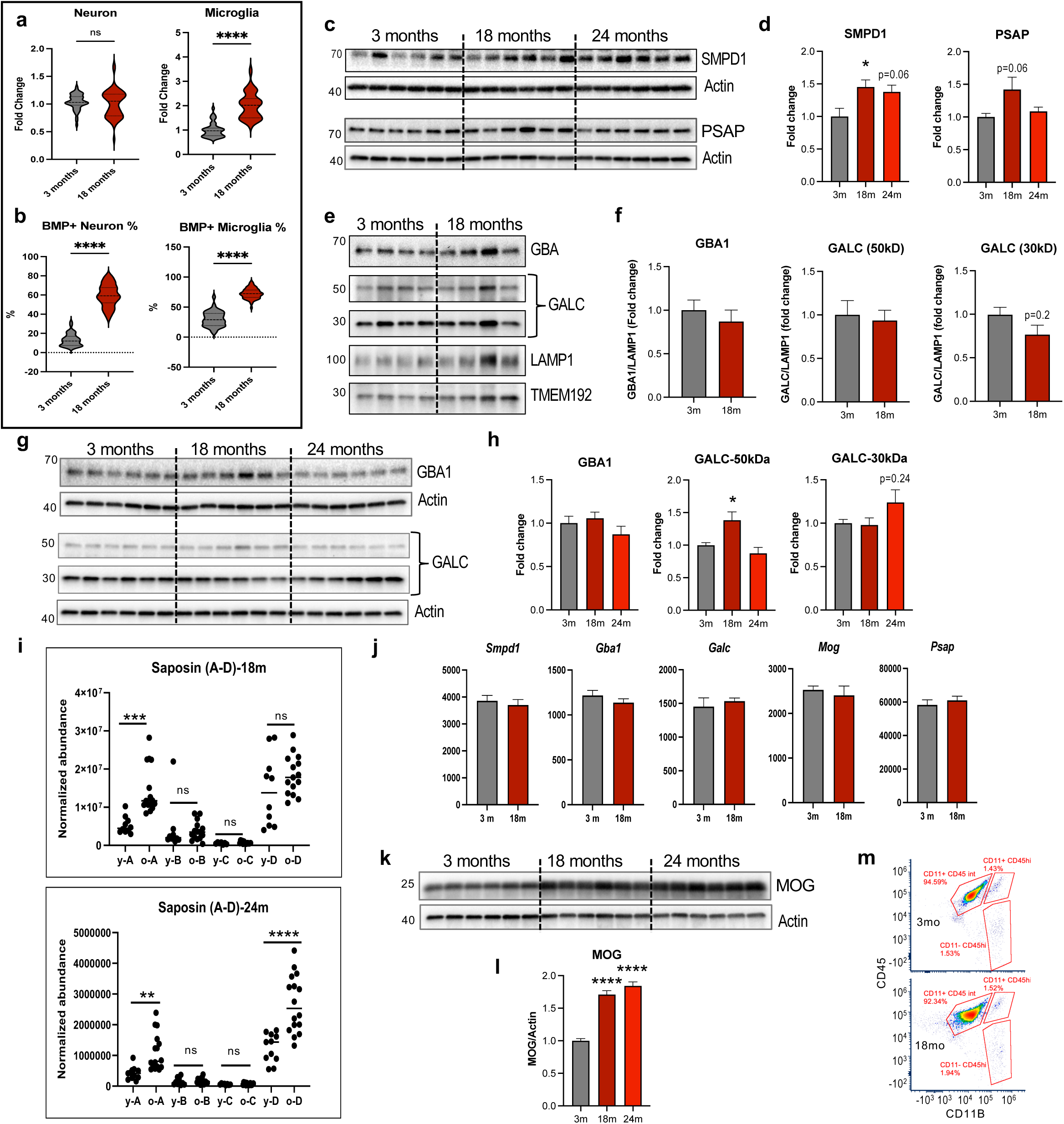
Lysosomal lipid metabolism is altered during brain aging. **a-b,** Quantification of data from Figure 6a-b, including total neuronal and microglial numbers (**a**) and fraction of corresponding cells positive for BMP (**b**). \*\*\*\**p*-value < 0.0001, unpaired *t*-test with Welch’s correction; *n* = 4-5. **c-d,** Western blot and quantification of lysosomal SMPD1 and PSAP in the cortical tissue. **e-h,** Western blot and quantification of HexCer catabolic enzymes GBA1 and GALC (30 kDa and 50 kDa fragments) in purified lysosomes (e-f) and total cortical lysates (g-h). **i,** MS analyses of lysosomal abundance of PSAP fragments, saposins A through D at 18 and 24 months. **j,** Tissue levels of corresponding mRNAs expression based on bulk RNA-seq analyses. **k-l,** Western blot and quantification of myelin component protein MOG in mouse cortical tissues. Data are mean +/- SEM; \**p*-value < 0.05; one-way ANOVA (b,f), Mann Whitney test (d) or unpaired t-test (g); n = 4-6. **m,** Flow cytometry strategy for identification of microglial cells in young (3 months) and aged (18 months) cortices based on CD11B and CD45 expression.

**Figure S7.**
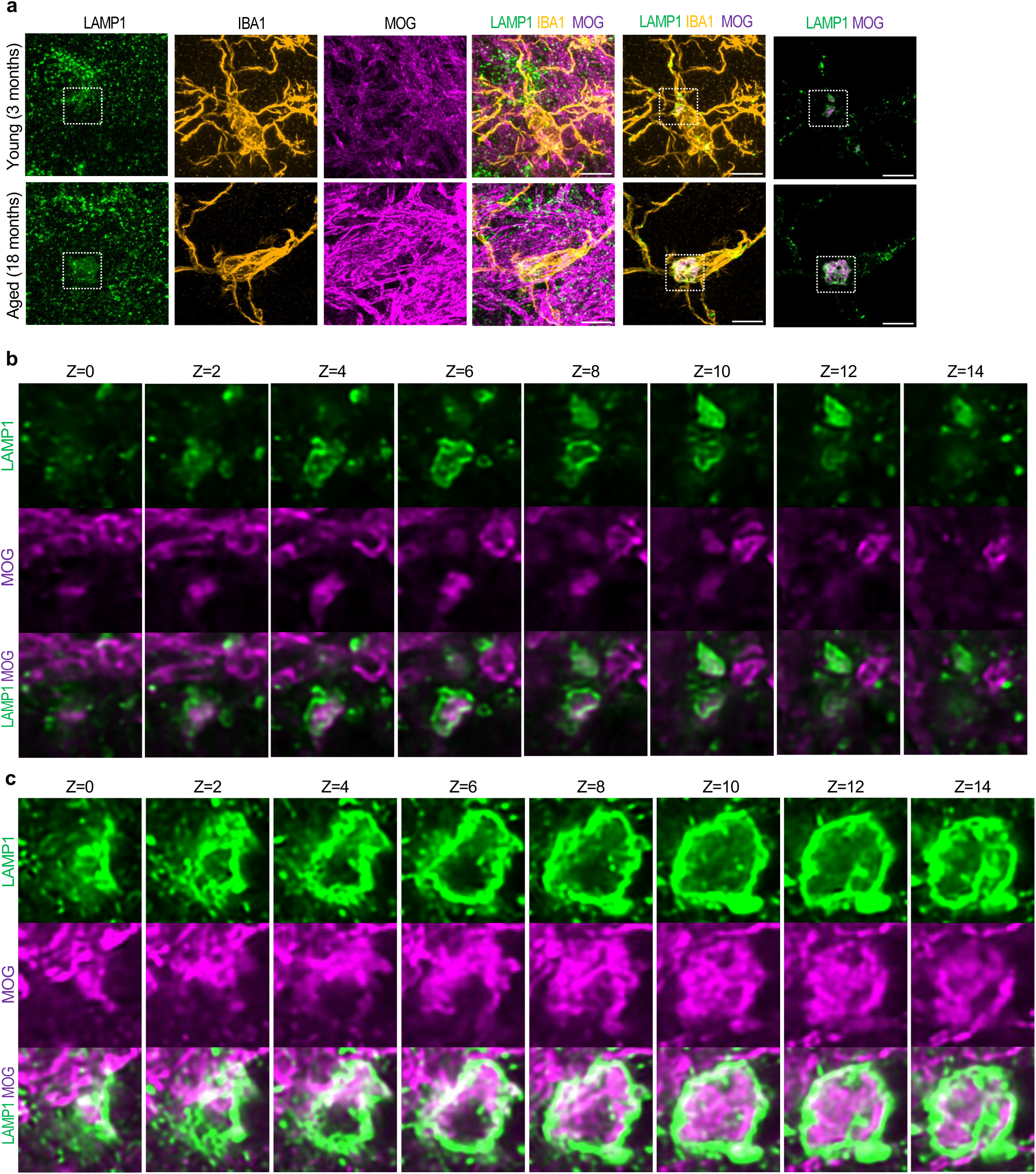
Quantification of lysosomal myelin accumulation during brain aging – related to Figure 6l-n. **a,** Single channel images and steps for quantification of lysosomal MOG in young and aged microglia. **b-c,** Closeup images of sequential z-stacks demonstrating localization of MOG within LAMP1 lysosomes in young (b) and aged (c) microglial cells.

**Figure S8.**
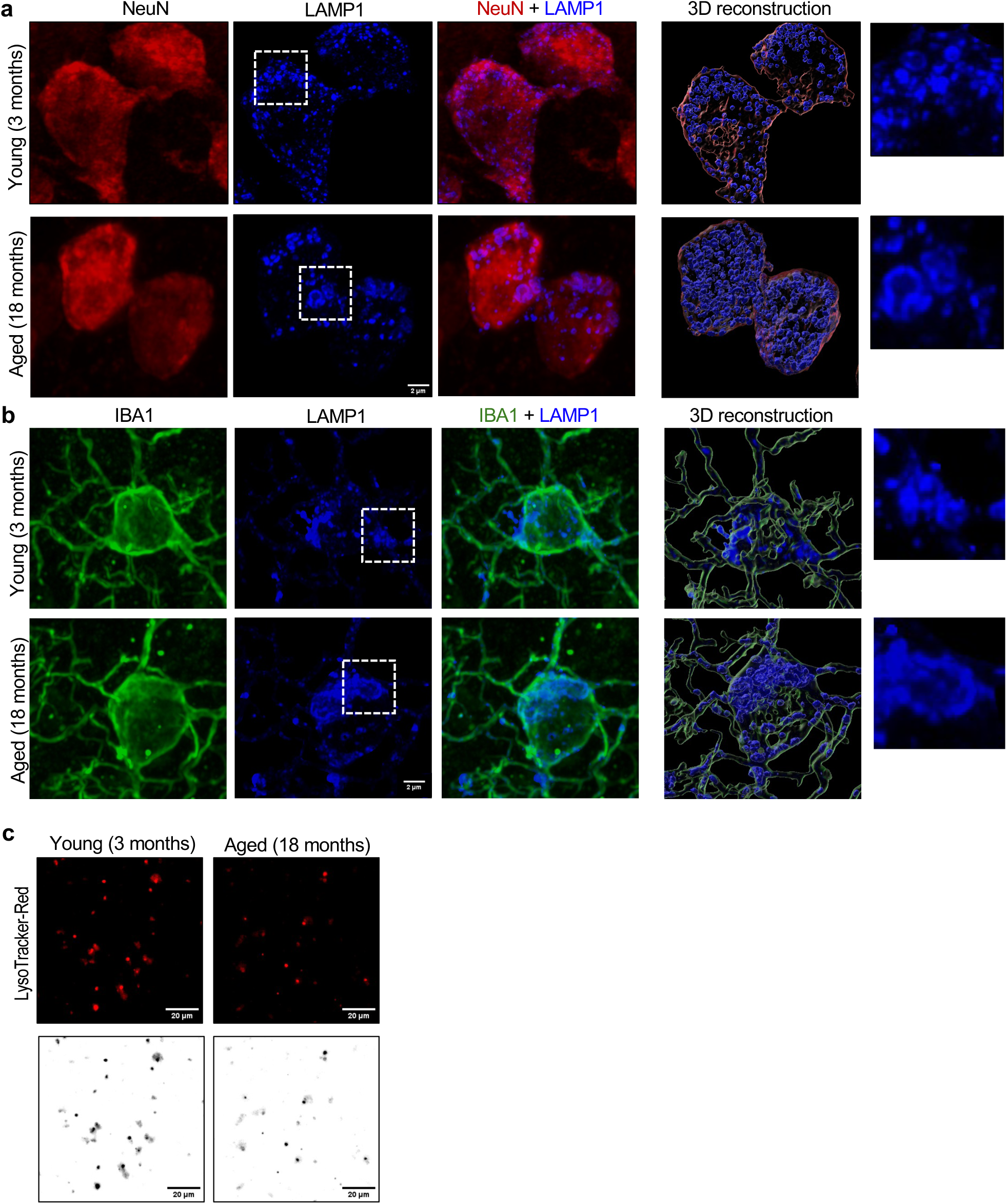
Quantification of lysosomal size and function in young and aged brain – related to Figure 7. **a-b,** Single channel images and steps in 3-D reconstruction used for quantification of lysosomal volume in young and aged neurons (a) and microglia (b). **c,** Images demonstrating altered LysoTracker intensity in lysosomes isolated from young as compared to aged mouse cortex.

**Figure S9.**
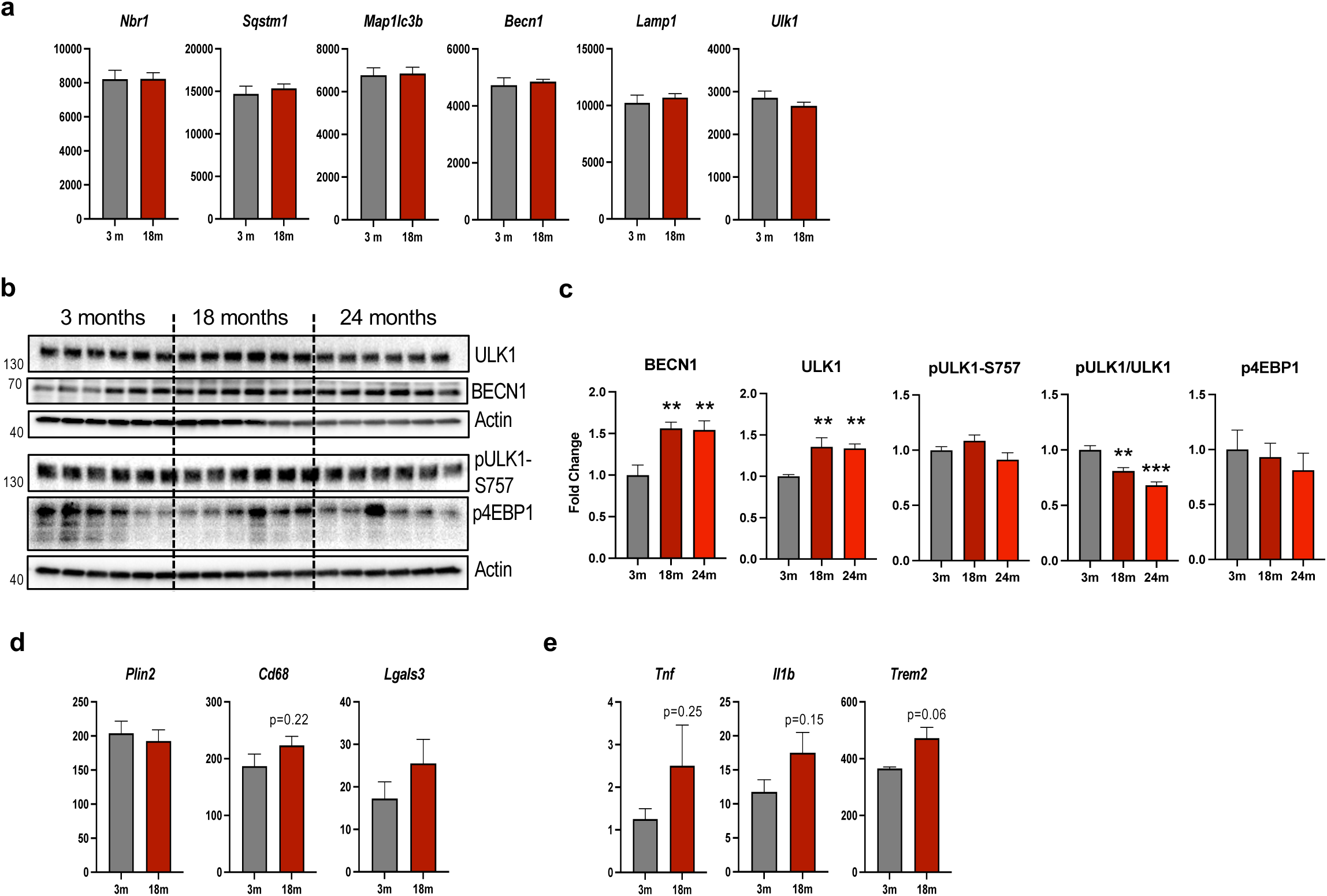
Quantification of autophagy and inflammation in young and aged brain. **a,** Bulk (mouse cortex) RNA-seq based quantification of autophagy and lysosomal gene expression in young and aged mouse cortex. **b-c,** Western blot and corresponding quantification of autophagy regulators BECN1 and ULK1 and phosphorylation of mTOR1 targets pULK1 and p4EBP1. **d,** Quantification of corresponding gene expression changes. **e,** Quantification of age-related gene expression changes of inflammatory genes. Data are mean +/- SEM; *I-value < 0.05; **I-value < 0.01; unpaired t-test or t-test with Welch’s correction (a, d-e) or 1-way ANOVA (c); n = 4-6

## Methods

### Animals

All experiments were performed according to NIH guidelines for the care and use of animals in research and under protocol approved by IACUC at the University of Maryland, Baltimore. All animals were males on *C57Bl6/J* background (Jackson Laboratory, Bar Harbor, ME), housed on sterilized bedding in a specific pathogen-free facility (12-hour light/dark cycle). Mice were aged in house to 12, 18 or 24 months prior to euthanasia for cortical sample collection (*N* = 8). Each hemisphere was processed separately. To minimize the technical variation, separate groups of samples from 3-months-old animals were collected and processed with each aged group (*N* = 5 or 6). Additional controls included identical pooled samples from 3-month-old animals, which were included with each of the age groups (*N* = 3). Inclusion of these controls allowed us to identify sources of any observed batch effects (sample collection versus MS instrument run) and to apply robust computational methods for batch correction (described below). Animals in 18- and 24-months groups plus 3-month-old controls underwent behavioral assessments prior to sample collection.

### Neurobehavioral assessments

Beam Walk: Fine motor coordination was assessed using the beam walk test as previously described. ^59^ Briefly, mice were placed at the end of a wooden beam (5 mm wide, 120 mm long) and the number of slips (foot faults) of the right hind limb were recorded over 50 steps. Mice were assessed by beam walk for 4 consecutive days for learning curve calculations at 3, 18 and 14 months old (*N* = 9-18).

Novel Object Recognition (NOR): Non-spatial memory in mice was assessed by performing NOR testing as previously described^59^. On test day, mice were allowed to freely explore for until 20 s of total interaction between the objects was recorded. Novel/familiar objects and side in which the novel object was positioned were balanced across all experimental groups to control for any potential object and side biases. Testing was recorded using Any-Maze software. Novel object preference was quantified as the ratio of time spent with the novel object to the time spent with the familiar object. Mice were tested at 3, 12, 18 and 24 months old (*N* = 14-28).

Morris Water Maze (MWM): Hippocampus-dependent spatial learning and memory were assessed using the MWM as previously described^59^. A circular tank (100 cm in diameter) was filled with water (24 ± 2°C) and was surrounded by various extra-maze on the inner walls of the testing area. A transparent platform (10 cm in diameter) was submerged 0.5 cm below the surface of the water. Mice were trained to find the hidden submerged platform located in the northeast (NE) quadrant of the tank for 4 consecutive days. The mice underwent four trials per day, starting from a randomly selected release point (east, south, west, and north). Each mouse was allowed a maximum of 90 s to find the hidden submerged platform. Testing was recorded using Any-Maze software. To find the slope of the learning curve the trial times for each day were averaged. Search strategy analysis was performed, characterized as follows: 1) Spatial search strategy: swimming directly to platform with no more than one loop or swimming directly to the correct target quadrant and searching for the platform; 2) Sequential search strategy: searching the interior portion of or the entire tank without spatial bias, including searching within the incorrect target quadrant before finding platform; and 3) Looping search strategy: circular swimming around the tank, swimming in tight circles, or swimming around the wall of tank before finding the submerged hidden platform. Mice were tested at 3, 12, 18 and 24 months old (*N* = 8-18).

### Western blots

Mice were perfused with saline, and one cortical hemisphere was dissected and homogenized in RIPA buffer (TekNova, Hollister, CA, USA; Cat. No. R3792) supplemented with protease inhibitors (Roche, Cat. No. 11836170001) and phosphatase inhibitors (Sigma, Cat. No. P5792), as previously^77^. Approximately 10-15 µg of total protein from tissue lysates was separated on 4–20% SDS-PAGE gel (Bio-Rad Laboratories, Cat. No. 5671095) and transferred to PVDF membranes (Millipore Sigma, Cat. No. IPVH00010) using a semi-dry transfer system. Membranes were blocked with 5% nonfat milk in 1× TBS containing 0.05% Tween-20 (TBST), then incubated overnight at 4 °C with primary antibodies diluted in 1% BSA in TBST. After washing, membranes were incubated for 1 h at room temperature with HRP-conjugated secondary antibodies diluted in blocking buffer. Protein bands were visualized using chemiluminescent substrates SuperSignal West Dura (Thermo Fisher Scientific, Cat. No. 34076), SuperSignal Femto (Cat. No. 34080), SuperSignal Atto, or SuperSignal West Pico (Cat. No. 34080) and imaged using the ChemiDoc Universal Hood II system (Bio-Rad Laboratories). Band intensities were quantified using Image Lab software (Bio-Rad Laboratories) and normalized to appropriate loading controls.

All lysates were resolved on 4–20% SDS-PAGE gels (Bio-Rad Laboratories, 5671095) and transferred to PVDF membrane (Millipore Sigma, IPVH00010) using semi-dry transfer. Membranes were blocked with 5% nonfat milk, probed with primary antibodies overnight at 4°C and incubated with HRP-conjugated secondary antibodies (Sera-Care KPL, 474–1506, 474– 1806, 14–16–06 and 14–13–06) in blocking solution at room temperature for 1 h. Protein bands were detected using SuperSignal West Dura Extended Duration Substrate (Thermo Fisher Scientific, 34076), SuperSignal Femto Chemiluminescent Substrate (Thermo Fisher Scientific, 34080) or SuperSignal West Pico Chemiluminescent Substrate (Thermo Fisher Scientific, 34080) and visualized using Chemi-doc system (Bio-Rad Laboratories, Universal Hood II). Band intensity was analyzed using Image Lab software (Bio-Rad Laboratories) and normalized to loading control. Antibodies used in this study are listed in Table S20 below.

### Lysosomal purification

Lysosomal fractions were isolated from the cortical brain tissues of mice using Lysosomal purification kit (Thermo Fisher Scientific, 89839) following the manufacturer’s instruction^16^.

### Lysosomal activity assays

CTSD activity was determined using fluorometric cathepsin D assay kit (Biovision Inc, K143-100) following manufacturers’ protocols. Enzyme activity in the lysosomal fraction and whole tissue lysate was quantified as change in fluorescence per μg of protein.

LysoTracker staining: Isolated lysosomes were resuspended in 50 µl of 1XPBS. Suspension (15 µl) was deposited onto glass coverslip coated with 100 µg/mL Poly-L-Lysine(Sigma-Aldrich, P6282), washed 3 times with PBS and incubated with 100 nM LysoTracker (Invitrogen, L7528) for 15 min. Coverslips were washed 3 times in PBS before being mounted on glass slide and imaged. For quantification, images were acquired and analyzed and quantified using Elements software (V5.30.06, Nikon) on a fluorescence Nikon Eclipse Ni-E, using a 20x/0.75 Plan-Apochromat Lambda DIC objective lens (Nikon). Briefly, cells positive for IF markers were identified using Detect Regional Maxima algorithm, followed by global threshold adjustment.

### Immunofluorescence staining

Anesthetized animals were transcardially perfused with cold saline, then with 4% paraformaldehyde (PFA, pH 7.4). Brains were postfixed with 4% PFA for 24 h, cryoprotected first in 20%, then in 30% sucrose for a total of 48 h and cut into 30 μm frozen sections. Sections were washed twice in PBS solution and incubated with 100mM Glycine (company) for 20 min at RT. Following 2 washes with PBS, tissues were incubated with permeabilization buffer (PB; 0.5% Triton X-100 in PBS) for 30 min and with blocking buffer (BB; 5% normal goat serum (Millipore, S26-100), or 5% normal donkey serum (Millipore, S30-100) in 0.1% Triton X-100 supplemented with 1% BSA) for 1 h, at RT. Samples were incubated with primary antibodies in BB at 4°C overnight. The following day, samples were washed 3 times with 1X PBS + 0.025% Triton X-100, before being incubated with secondary antibodies in the BB for 2 h, at RT. Nuclei were stained with 1 μg/ml Hoechst (Invitrogen, H1399). Sections were mounted on glass slide and coverslips using Hydromount solution (National Diagnostics, 50–899-90,144). For BMP staining, 0.2% saponin was used in placed of Triton X-100 for PB and BB. For STED, BB (2% BSA, 0.1% Triton X-100) was used. Secondary antibodies were incubated for 3 hr. Following secondary incubation, sections were washed 3 times with 1X PBS and coverslipped using Abberior MOUNT liquid antifade solution (Abberior, MM-2009). Antibodies are listed in Table S20 below.

### Wide field image acquisition and analysis

Images were acquired on a fluorescence Nikon Eclipse Ni-E microscope using Nikon 20x/0.75 Plan-Apochromat Lambda DIC objective lens (BMP) or 60x/1.40 Plan-Apochromat Lambda DIC objective lens (MOG). Emission wavelengths included DAPI (420-460 nm), FITC (515-555 nm), TRITC (577-630 nm), Cy5 (677-711 nm) and Cy7 (768-849 nm). Exposure times were kept constant for all sections within the experiment. BMP analysis: z-stacks were captured at 1 µm intervals and processed with the NIS-Elements Extended Depth of Focus (EDF) algorithm to reconstruct a single in-focus composite image. Images were analyzed and quantified by Elements software (V5.30.06, Nikon) as described: nuclei were identified using Spot Detection algorithm; cells positive for IF markers were identified using Detect Regional Maxima algorithm, followed by global threshold adjustment^59^. MOG analysis: Z-stacks were acquired at 0.3 μm intervals across a total depth of ∼10 μm to ensure full cell coverage. To enhance signal-to-noise ratio and spatial resolution, raw images were processed using the NIS-Elements Denoising and Deconvolution Module (Nikon Instrument). Images were analyzed for mean fluorescence intensity (MFI) of MOG inside of IBA1-positive LAMP1 signals. Cells positive for IF markers were identified using Rolling Ball algorithm, followed by global threshold adjustment. Only thresholds that are positive for both IBA1 and LAMP1 were kept, LAMP1 and MOG MFI were measured. Data were normalized to the respective IBA1 volumes.

For representative images, confocal images were acquired using the Nikon CSU-W1 microscope equipped with 405, 488, 561, and 647 lasers, using a 60x (1.49 NA) TIRF oil-immersion and Nikon Elements software. Microglial surfaces of fluorescence images were reconstructed using the ‘surface’ feature and default parameters. Following, all the lysosomal signal outside of the reconstructed surfaces were masked to ‘0’, retaining only lysosomal signal inside the reconstructed surfaces.

### Airyscan image acquisition and analysis of lysosomal morphology

Images were acquired using a Zeiss LSM 880 microscope (Zeiss Microimaging) equipped with an Airyscan superresolution imaging module, using a 63x/1.40 Plan-Apochromat Oil DIC M27 objective lens (Zeiss Microimaging). The 488-nm Argon laser line, 561-nm DPSS 561 laser, and 633-nm HeNe 633 laser were used to detect Alexa-488, Alexa-568, and Alexa-633, respectively. Acquisition parameters were kept constant. For quantification, z-stacks covering the entire depth of the cells with steps of 0.18 µm were acquired, followed by the Airyscan image processing, set at medium strength. For representative images, the images were processed using Airyscan Joint Deconvolution (jDCV) method, resulting per Zeiss Microimaging in generation of ∼90 nm lateral resolution images^78^.

Quantification was performed using Imaris Bitplane software (10.2, Oxford Instrument). Briefly, neuronal and microglial surface of fluorescence images were reconstructed using the ‘surface’ feature and default parameters. Following, all the lysosomal signal outside of the reconstructed surfaces were masked to ‘0’, retaining only lysosomal signal inside the reconstructed surfaces. After masking, lysosomal surfaces were reconstructed using the ‘surface’ feature and ‘machine learning’ algorithm parameters. Reconstructed lysosomes with volumes greater than 0.5 μm^3^ were counted as enlarged.

### Stimulated emission depletion (STED) imaging and analysis

Images were acquired with the Abberior Facility Line STED microscope at the Confocal Microscopy Core Facility at UMB. STED imaging was performed using an Olympus 60x/1.42 STED-UPLXAPO 60 oil immersion lens. Abberior STAR Orange and STAR Red were excited at 561 nm and 640 nm respectively and depletion was performed with a 775 nm pulsed laser with a gating of 1-7 ns and a pixel dwell time of 5 µs. The emission from both channels was collected with the matrix detector; images were post-processed with the Abberior Imspector software. Each line was scanned 15 times during acquisition (line accumulations). The three-dimensional STED data was acquired using the Easy3D module with voxel sizes set to 30 nm and the pinhole was set to 1 AU.

### Flow cytometry

Cortices were removed and processed by mechanical disruption on a 70 μm filter screen and resuspended in RPMI-1640 (Quality Biological, 112-025-101). Ten Units (U) Papain (Sigma-Aldrich, P4762), 10 mg/ml DNase II (Sigma, 10104159001), 1 mg/ml Collagenase-Dispase (Sigma, 10269638001), and 1 μl of GolgiPlug containing brefeldin A (BD Biosciences, 555029) were added to the brain suspension and incubated on a shaker at 200 rotations per minute (RPM) for 1 hour at 37 °C for mechanical and enzymatic digestion of the brain tissue. After incubation, leukocytes were separated from other brain cells by a Percoll gradient (GE Healthcare, Chicago, IL, USA, GE17-5445-02). Brain leukocytes were resuspended in 70% Percoll-HBSS and were slowly injected under a 30% Percoll-RPMI layer using a blunt popper pipetting needle (Sigma). This Percoll gradient was spun for 20 min with no brake. Myelin layer was removed from the top of the 30% Percoll layer and leukocytes were then retrieved from the interface of the 30% and 70% Percoll layers and resuspended in RPMI as single cell suspensions. These leukocytes were then washed in fluorescence-activated cell sorter (FACS) buffer (5% FBS and 0.1% penicillin and streptomycin in 1xHBSS) with sodium azide and blocked with 1:50 mouse Fc Block (clone 93; eBioscience, San Diego, CA, 14-0161-82) for 10 min on ice prior to staining with primary antibody-conjugated fluorophores. After primary antibody staining, cells were washed in Fixation/Permeabilization solution (BD Biosciences, 554722) for 20 min and washed twice in Permeabilization/Wash Buffer (BD Biosciences, 554723) and resuspended in an intracellular antibody cocktail of cytokine antibodies. After intracellular antibody staining for 30 min at 4 °C, samples were washed with Wash Buffer, fixed with 2% PFA and resuspended in FACS buffer. Data were acquired on a BD LSRII flow cytometer equipped with FACsDiva 6.0 (BD Biosciences, San Jose, CA) and analyzed using FCS ExpressTM 7 (De Novo Software, Glendale, CA). Antibodies are listed in Table S20 below.

### Statistical analyses for follow-up experiments

Analyses were performed using GraphPad Prism Software v. 9.2 (GraphPad Software, Inc., La Jolla, CA), using one-way ANOVA with Tukey’s or Dunnett’s multiple comparison test. Experiments with two groups were analyzed using unpaired two-tailed *t*-test (groups with equal variance) or *t*-test with Welch’s correction (unequal variance). Beam walk data was analyzed using 2-way repeat measures (RM) ANOVA with Sidak’s post-hoc and age and time (learning days) as variables. Statistical analyses and sample numbers for each experiment are specified in figure legends. Unless otherwise specified all *P* values reflect comparisons with young (3-months-old) mice. *P* < 0.05 was considered significant. Bar graphs are mean ± SEM; for behavioral experiments dots correspond to individual animal values; dotted lines in violin plots indicate median and top/bottom quartiles.

### Mass spectrometry lipidomics and analyses

LC-MS grade acetonitrile, methanol, water, and n-propanol were purchased from Fisher Scientific (Pittsburg, PA). HPLC grade tert-Butyl methyl ether (MTBE), chloroform, ammonium formate, and formic acid was purchased from Sigma Aldrich (St. Louis, MO). EquiSPLASH lipidomix was purchased from Avanti Polar Lipids, Inc. (Alabaster, AL). Total lipid extracts were prepared using a modified MTBE lipid extraction protocol^33,79^. Briefly, 400 µL of cold methanol and 10 µL of internal standard mixture (EquiSPLASH lipidomix) were added to each sample followed by incubation at 4°C, 650 rpm shaking for 15 minutes. Cold MTBE (500 µL) was added followed by incubation at 4°C for 1 hour with 650 rpm shaking. Cold water (500 µL) was added slowly and resulting extract was maintained 4°C, 650 rpm shaking for 15 minutes. Phase separation was completed by centrifugation at 8,000 rpm for 8 min at 4 °C. The upper, organic phase was removed and set aside on ice. The bottom, aqueous phase was re-extracted with 200 µL of MTBE followed by 15 minutes of incubation at 4°C with 650 RPM shaking. Phase separation was completed by centrifugation at 8,000 rpm for 8 min at 4 °C. The upper, organic phase was removed and combined with previous organic extract. The organic extract was dried under a steady stream of nitrogen at 30 °C. The recovered lipids were reconstituted in 200 µL of chloroform:methanol (1:1, v/v) containing 200 µM of butylated hydroxytoluene. Prior to analysis samples were further diluted 10-fold with acetonitrile:isopropanol:water (1:2:1, v/v/v). The lower aqueous phase was used to determine the protein content via a BCA kit (bicinchoninic acid assay, Thermo Fisher Scientific, Rockford, USA).

Total lipid extracts were analyzed by liquid chromatography coupled to high-resolution tandem mass spectrometry (LC-MS/MS)^33,34^. The LC-MS/MS analyses were performed on an Agilent 1290 Infinity LC coupled to an Agilent 6560 Quadrupole Time-of-Flight (Q-TOF) mass spectrometer. The separation was achieved using an C18 CSH (1.7 µm; 2.1 × 100 mm) column (Waters, Milford, MA). Mobile phase A was 10 mM ammonium formate with 0.1% formic acid in water/acetonitrile (40:60, v/v) and mobile phase B was 10 mM ammonium formate with 0.1% formic acid in acetonitrile/isopropanol (10:90, v/v). The gradient was ramped from 40% to 43% B in 1 min, ramped to 50% in 0.1 min, ramped to 54% B in 4.9 minutes, ramped to 70% in 0.1 min, and ramped to 99% B in 2.9 min. The gradient was returned to initial conditions in 0.5 min and held for 1.6 min for column equilibration. The flow rate was 0.4 mL/min. The column was maintained at 55 °C and the auto-sampler was kept at 5 °C. A 2 µL injection was used for all samples. Mass spectrometry analysis was separated into two workflows: 1) lipid identification of a pooled sample using an iterative MS/MS acquisition and 2) lipid semi-quantitation of all samples using high-resolution, accurate mass MS^1^ acquisition. The MS parameters for the iterative workflow were as follows: extended dynamic range, 2 GHz; gas temperature, 200 °C; gas flow, 10 L/min; nebulizer, 50 psi; sheath gas temperature, 300 °C; sheath gas flow, 12 L/min; VCap, 3.5kV (+), 3.0kV (–); nozzle voltage, 250V; reference mass *m/z* 121.0509, *m/z* 1221.9906 (+), *m/z* 119.0363, *m/z* 980.0164 (–); MS and MS/MS Range *m/z* 100–1700; acquisition rate, 3 spectra/s; isolation, narrow (∼ 1.3 *m/z*); collision energy 20 eV (+), 25 eV (–); max precursors/cycle, 3; precursor abundance-based scan speed, 25,000 counts/spectrum; ms/ms threshold, 5,000 counts and 0.001%; active exclusion enabled yes; purity, stringency 70%, cut off 0%; isotope model, common organic molecules; static exclusion ranges, *m/z* 40 to 151 (+,–). The MS parameters for the MS^1^ workflow were the same for source and reference mass parameters and differed only for acquisition (selection of MS (same parameters) not Auto MS/MS).

LC-MS data from the iterative MS/MS workflow was analyzed for lipid identification via Agilent’s Lipid Annotator (v 1.0). The default settings for feature finding and identification parameters were used. Positive and negative ion mode adducts included [M+H]^+^, [M+Na]^+^, [M+NH_4_]^+^, [M-H]^−^, and [M+CH_3_CO_2_]^−^, respectively. The results of the Lipid Annotator were saved as a PCDL file. The LC-MS data from the MS^1^ workflow were processed using Agilent’s MassHunter Profinder (v 10.0). Batch targeted feature extraction using default parameters and the PCDL file created from Lipid Annotator were used for feature extraction. The processed data generated from Profinder which included peak area and lipid identification was exported into MetaboAnalyst 5.0^80^ for multivariate analysis. Univariate analysis was done using Prism 6 (GraphPad, La Jolla, CA).

### Mass spectrometry proteomics and analyses

Lysosomal samples were collected and stored at −80 °C until analysis. Protein extraction and digestion were performed using Pierce™ Mass Spec Sample Prep Kit for Cultured Cells (Thermo Fisher Scientific, Cat. No. 84840) according to the manufacturer’s instructions. Briefly, lysates were prepared by adding five volumes of lysis buffer to the cell pellet, followed by washing, reduction, alkylation, and enzymatic digestion with trypsin. Digested peptides were quantified using a BCA assay kit (Thermo Fisher Scientific, Cat. No. 23275) to determine peptide concentration. LC-MS/MS-based proteomic analysis was conducted on a nanoACQUITY Ultra-Performance Liquid Chromatography system (Waters Corporation, Milford, MA USA) coupled to an Orbitrap Fusion Lumos Tribrid mass spectrometer (Thermo Scientific, San Jose, CA USA). Peptide separation was effected on a nanoACQUITY Ultra-Performance Liquid Chromatography (UPLC) analytical column (BEH130 C18, 1.7 µm, 75 µm x 200 mm; Waters Corporation, Milford, MA, USA) using a 185-min linear gradient with 3-40% acetonitrile and 0.1% formic acid. Mass spectrometry conditions were as follows: full MS scan resolution of 240,000, precursor ions fragmentation by high-energy collisional dissociation of 35%, and a maximum cycle time of 3 s. The Pierce HeLa Protein Digest Standard (Thermo Fisher Scientific, #88329) was injected between runs as an instrument quality control to monitor system performance. The resulting mass spectra were processed using Thermo Proteome Discoverer (PD, version 2.5.0.400, Thermo Fisher Scientific) and searched against a UniProt mouse reference proteome (release 2022.06, 17180 entries) using Sequest HT algorithm. Search parameters include carbamidomethylation of cysteines (+57.021 Da) as a static modification, methionine oxidation (+15.995 Da) as a dynamic modification, precursor mass tolerance of 20 ppm, fragment mass tolerance of 0.5 Da, and trypsin as a digestion enzyme. Tryptic missed cleavages were restricted to a maximum of two, with peptide lengths set between 6 and 144 residues.

For protein quantification, the Minora feature detector, integrated in the PD, was used as described previously^81^. To ensure high data quality, proteins were further filtered to a 1% false discovery rate (FDR) threshold, calculated with the Percolator algorithm. Next, protein abundance values exported from PD were post-processed using Perseus software (version 1.6.14.0). Proteins with missing values were excluded to improve data quality. The quantitative protein data were log2 transformed and further normalized using median centering. Overall, 4023 protein groups were identified at 12 months, 3463 at 18 months, and 2898 at 24 months, using 1% FDR threshold. Of these, 2986, 2004, and 2287 protein groups were quantified without missing values across all replicate samples at 12, 18, and 24 months, respectively. Differentially expressed proteins (DEPs) were identified using a two-tailed Student’s t-test (adjusted *P* < 0.05). Metabolanalyst (version 5.0) was used to generate heatmaps for comparison of aged groups with young controls.

### RNA sequencing

Total RNA was isolated from the left cortex of 3-, 12- and 18-month-old mice using Trizol reagent (Invitrogen, 15596–018) as per the manufacturer’s instruction. RNA sequencing and analysis was carried out by the Maryland Genomics, Institute for Genome Sciences, UMSOM. To minimize ribosomal RNA background, samples were enriched for mRNA containing Poly A tails using bead-based enrichment kit (NEBNext Poly(A) mRNA Magnetic Isolation Module, New England Biolabs). Enriched Poly A RNA was used to create Strand-specific, dual unique indexed libraries using the NEBNext Ultra II Directional RNA Library Prep Kit for Illumina (New England Biolabs). Manufacturer protocol was modified by diluting adapter 1:30 and using 3 ul of this dilution. Size selection of the library was performed using SPRI-select beads (Beckman Coulter Genomics). Glycosylase digestion of adapter and 2nd strand was done in the same reaction at the final amplification. Revvity GX touch was used to assess libraries for size and quantity. The libraries were pooled, assessed by qPCR using the KAPA Library Quantification Kit (Complete, Universal, Kapa Biosystems) and sequenced on an Illumina NovaSeq 6000 using 100bp PE reads. Paired-end Illumina libraries were mapped to the Mouse reference, Ensembl release GRCm39.108, using HiSat2 v2.0.4, using default mismatch parameters. Read counts for each annotated gene were calculated using HTSeq. The DESeq2 Bioconductor package (v1.5.24) was used to estimate dispersion, normalize read counts by library size to generate the counts per million for each gene, and determine differentially expressed genes between disease and control samples. Differentially expressed transcripts with FDR ≤ 0.05 and log2 fold change ≥ 2 were used for downstream analyses.

### Data preparation for machine learning analyses

Mass spectrometry data were compiled into six Microsoft Excel spreadsheet files, with each file corresponding to a different -omics type (lipid or protein) or age group (12-month-old, 18-month-old, 24-month-old). Control mice (3-month-old) were included with each file. Protein mass spectrometry data were collected in six batches, corresponding to brain hemisphere and non-control age groups (with control mice in each batch). Lipid mass spectrometry data were collected in three batches, corresponding to three non-control age groups. Protein data were formatted with proteins (Uniprot accession IDs ^82^) as rows and samples as columns and subsequently transposed. Lipid data were formatted with samples as rows and lipids species as columns. Lipid species names were standardized to LIPIDMAPS shorthand hierarchical nomenclature. ^83^ Where multiple isomers were detected and more specific structure was not determined, subscripts (e.g., “_i1”, “_i2”) were appended to the name to distinguish isoforms. All data analytical work was performed in R. ^84^

Robust principal component analysis (PCA) ^85^ implemented in the rospca R package ^86^ was used to identify outliers. Outliers were detected individually for each batch, using an α cutoff of 0.95. Outliers flagged by either score distance or orthogonal distance were removed prior to batch correction and from subsequent analyses.

Batch correction was conducted using a location-and-scale model. ^87^ We first examined whether our mass spectrometry abundance data fit a log-normal, exponential, or Weibull distribution (non-negative continuous distributions). Each group of technical and biological replicates was ranked for each distribution by log-likelihood, with fit to a log-normal distribution generally being the best. The data were then assumed to have the location-and-scale model form (modified from previous work ^87^)

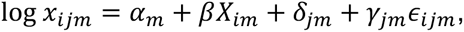

where *i* is the sample, *j* represents the batch, *m* is the molecule (i.e., a protein or lipid species), *⍺* is the abundance over all samples, *X* is the set of experimental conditions, *β* is the change in abundance corresponding to a set of experimental conditions, *δ* is a location batch effect, *γ* is a scaling batch effect, and *ε* is a normally distributed random error. In our experimental design, a single set of biological conditions were shared across batches and used as controls (3-month), therefore we estimated batch effects using these samples. For each batch, the location for each batch was estimated using the trimmed mean, removing 20% of the highest and lowest values, which made the estimation more robust to outliers, and the scale was estimated using the standard deviation. The true location and scale were assumed to be the median location and scale across batches, and each batch was adjusted to the true location and scale.

Batch effects were assessed before and after batch correction by three methods: 1) visually with the major principal components after removing lipid species with missing values; 2) applying random forest ^88^ classification to abundance values; and 3) applying random forest classification to principal component scores for the components that explain the top 95% of variance. Models were constructed with the batch as the response, comprised 10,000 trees (50,000 trees when using protein abundance values, given the greater number of predictors), and larger classes were down-sampled to address imbalanced class sizes. Models assessing batch effects were first constructed on all samples but evaluated on biological conditions that spanned both batches. A one-sided binomial test of model performance, implemented in the ‘confusionMatrix’ function of the caret R package, ^89^ was used to obtain a *P* value comparing model accuracy to the no information rate.

### Multi-omics factor analysis and machine learning analyses

Multi-omics factor analysis (MOFA) ^90^ was conducted on log_10_ transformed data. Protein samples where corresponding lipid samples were removed as outliers or were missing were removed prior to analysis (as exploratory results showed that these were likely to yield outliers through MOFA imputation). Views were labeled as either “protein” or “lipid”. A combination spike-and-slab and ARD sparsity prior was used for the factors, and a spike-and-slab prior was used for the weights. Ten factors were estimated from this data set. MOFA models used for downstream analysis were evaluated for residual batch effects. This was done in the same manner as with principal component analysis and abundance data, using classification random forest with 10,000 trees and assessing model accuracy on control samples as a measurement of batch effects.

Factors were considered associated with a given biological covariate of interest if they were in the feature set determined by forward feature selection. Multiple testing for linear model coefficients was addressed by controlling the false discovery rate using the Benjamini-Hochberg method ^91^ across all linear models. Factors were considered to be associated with an -omics type if they explained greater than 1% of the variance in that -omics type. Factors associated with lipid variance were characterized by Lipid Ontology (LION) enrichment analysis, ^92^ and factors associated with protein variance were characterized using Gene Ontology, ^93^ transcriptional regulatory relationships (TRRUST), ^94^ and protein interaction enrichment analysis. ^95^

Forward feature selection was conducted to determine factors most closely associated with age. This was done using a forward feature selection ML framework, ^96,97^ where the initial feature ranking was determined by permutation feature importance determined by a regression random forest built on all features. ^98^ The feature list was first filtered using the elbow method as described below, and regression random forests were constructed using iteratively larger feature sets using this feature list. Models and associated feature sets were compared using their Akaike Information Criterion (AIC) ^99^ which was calculated as AIC = *n* log MSE + 2*k*, where *n* is the number of samples, MSE is the mean squared error, and *k* is the number of features used in each model. Factors yielding the model with the lowest AIC were considered to be associated with aging.

The elbow rule heuristic was used to determine a discrete cutoff based on a univariate measure in several analyses. This was used both for deciding which factors were associated with aging and for downstream analyses of factors. To calculate the elbow method cutoff, values were ordered paired with their rank. The rank-value pair yielding the largest absolute diagonal distance from the line calculated between the lowest and highest rank-value pairs was taken to be the elbow point.

### Enrichment and ontology analyses

We used the lipid ontology (LION) ^92^ for lipid enrichment analysis using the LION web tool at http://lipidontology.com/ with analysis parameters “from high to low” (indicating that higher values are positively associated with the factor) and “two-tailed”. Mappings between input lipid IDs and LION IDs were manually checked and missing mappings were added. False discovery rate was controlled using the Benjamini-Hochberg method. Four terms: CAT0000000, lipid classification; CAT:0000091, physical or chemical properties; CAT:0012007, function; and CAT:0012008, cellular component were used as overarching categories of ontology terms when examining terms, similar to the three aspects of Gene Ontology, Biological Process, Molecular Function, and Cellular Component.

We performed enrichment analysis using a modified version of the ‘run_enrichment’ function available in the ‘MOFA2’ R package, ^100^ which implements a modified version of principal component gene set enrichment. ^101^ MOFA2 R code was modified to accommodate parallel processing using the ‘parallel’ R package ^102^ via forking. The sign was set to “all” to capture both positive and negatively enriched term sets. The “rank-sum” test statistic was used with the “permutation” statistical test. Various term sets were generated from different databases as described in other methods.

Associations between proteins and GO terms were obtained from the ‘org.Mm.eg.db’ ^103^ and ‘org.Hs.eg.db’ ^104^ R packages. To obtain a fuller set of GO terms, mouse proteins were mapped to human orthologs using the NCBI datasets command line tool. ^105^ Any GO term associated with a mouse protein, or the corresponding human protein homolog was included as a GO annotation. These feature sets were then used as inputs as described for enrichment analysis.

Mouse transcription factor regulated gene lists were obtained from the TFLink gateway (https://tflink.net/download/ accessed on 4/8/2025). ^106^ The simple format gzipped file included UNIPROT Ids which were directly mapped to our data and were then converted into a binary protein set matrix compatible with the MOFA enrichment function.

Protein interaction information was retrieved from Uniprot ^82^ using the ‘UniprotR’ R package ^107^ including binary interactions and subunit structures. Because of the discrete binary nature of these data, enrichment analysis could not be used to test for enrichment of protein interactions. Instead, a discrete factor weight cutoff was determined using the elbow method, and proteins with an absolute factor weight above the cutoff were analyzed for pairwise interactions. Permutation tests with 1000 permutations were used to test for over-enrichment of protein interactions compared to randomly drawn sets of proteins, and the Benjamini-Hochberg method was used to control the false discovery rate.

We conducted over representation analysis (ORA) using the implementation from the ‘clusterProfiler’ R package. ^108^

Disease ontology enrichment analysis was performed using the over-representation analysis method as described above. DisGenNet ^109^ disease ontology information was obtained from the ‘DOSÈ R package, ^110^ and human Entrez gene ids were mapped to mouse Uniprot protein ids as described above using the ‘org.Mm.eg.db’ R package and the NCBI datasets command line tool. For each factor, the elbow method was used to identify a discrete set of proteins associated with the factor, which was then separated into the set of positively and negatively associated proteins for each factor before calculating disease ontology over-representation. The false discovery rate over all factor-disease associations was controlled using the Benjamini-Hochberg method.

### RNA-sequencing data analyses

RNA-seq initial data processing. Whereas proteomics and lipidomics data were collected according to a split sample study design, transcriptomics data (in relation to proteomics and lipidomics) were collected according to a repeated study design^111^, with four samples each of 3-, 12-, and 18-month-old mouse cortex. We normalized the total sample expression for each sample to the median total expression, and no outliers were removed.

To determine whether transcriptional patterns correlated well with proteomic abundance patterns, synthetic samples were generated by randomly pairing protein and RNA-seq samples matched by covariates (age), with the larger proteomics data set down-sampled to match the smaller RNA-seq data set for a total of twelve samples. Spearman rank correlation was then calculated between gene expression and protein abundance. Random pairings were repeated one hundred times, generating a 95% confidence interval (using the 2.5% and 97.5% quantiles) for the Spearman correlation value. Proteins with confidence intervals greater than zero were interpreted as correlated (i.e., the change with age in proteomics reflects that of transcriptomics). Proteins with confidence intervals less than zero were interpreted as anti-correlated, and proteins with confidence intervals overlapping zero were interpreted as having no definitive relationship between protein abundance and transcription. We conducted this analysis on the sets of proteins passing the elbow method cutoff for each of the aging-associated factors. Following identification of transcription-correlated proteins for each factor, we performed Gene Ontology over-representation analysis of transcription-correlated proteins using the full elbow method-cutoff set as the background.

**Table S20.** List of antibodies used in this study.

| Antibody | Procured from | Catalog No.<br># | Use | Dilution |
| --- | --- | --- | --- | --- |
| **Western blot – primary antibodies:** |  |  |  |  |
| MOG | R&D Systems | AF2439 | WB | 1:1000 |
| SMPD1 | Proteintech | 14609-1-AP | WB | 1:1000 |
| GALC | Proteintech | 11991-1-AP | WB | 1:1000 |
| ASAH1 | Proteintech | 11274-1-AP | WB | 1:1000 |
| GBA | Cell Signaling Technology | 88162 | WB | 1:1000 |
| PSAP | Proteintech | 10801-1-AP | WB | 1:1000 |
| Actin | Sigma | A1978 | WB | 1:5000 |
| LAMP1 | DSHB (deposited to the DSHB by August, J.T. of Johns Hopkins School of Medicine, Pharmacology & Molecular Sciences) | 1D4B | WB | 1:1000 |
| LAMP2 | DSHB (deposited to the DSHB by August, J.T. of Johns Hopkins School of Medicine, Pharmacology & Molecular Sciences) | ABL-93 | WB | 1:500 |
| TMEM192 | Proteintech | 28263-1-AP | WB | 1:1000 |
| **Secondary antibodies:** |  |  |  |  |
| Goat-anti-Rabbit-IgG-HRP-Conjugate | Sigma | 12-348 | WB | 1:2500 - 1:5000 |
| Goat-anti-Mouse-IgG-HRP-Conjugate | Sigma | 12-349 | WB | 1:2500 - 1:5000 |
| Goat-anti-Rat-IgG-HRP-linked Antibody | Cell Signaling Technology | 7077 | WB | 1:2500 |
| Goat-anti-Rabbit-IgG-HRP-linked Antibody | Cell Signaling Technology | 7074 | WB | 1:2500 |
| **Immunofluorescence – primary antibodies:** |  |  |  |  |
| IBA1 | Wako | 019-19,741 | IF | 1:200 |
| NeuN | MilliporeSigma | ABN90 | IF | 1:4000 |
| LAMP1 | DSHB | 1D4B | IF | 1:200 |
| LAMP1 | Abcam | ab24170 | IF | 1:200 |
| LBPA | Sigma-Aldrich | ZMS1107 | IF | 1:200 |
| TOM20 | Proteintech | PTG-11802 | IF | 1:200 |
| mMOG | R&D Systems | AF2439 | IF | 1:200 |
| **Secondary antibodies:** |  |  |  |  |
| Alexa Fluor 488 goat anti-mouse | Invitrogen | A11029 | IF | 1:200 |
| Alexa Fluor 488 goat anti-rat | Life Technologies | A11006 | IF | 1:200 |
| Alexa Fluor 568 goat anti-rabbit | Invitrogen | A11036 | IF | 1:200 |
| Alexa Fluor 633 goat anti-guinea pig | Invitrogen | A21105 | IF | 1:200 |
| Alexa Fluor 488 donkey anti-rat | Life Technologies | A21208 | IF | 1:200 |
| Alexa Fluor 568 donkey anti-goat | Invitrogen | A11057 | IF | 1:200 |
| Alexa Fluor 647 donkey anti-rabbit | Invitrogen | A31573 | IF | 1:200 |
| **STED antibodies:** |  |  |  |  |
| anti-mouse abberior STAR RED | Abberior | STRED-1001 | STED | 1:200 |
| anti-rabbit abberior STAR Orang | Abberior | STORANG E-1002 | STED | 1:200 |
| **Flow cytometry – surface antibodies:** |  |  |  |  |
| Zombie-Aqua | Biolegend | 423101 | FACS | 1:50 |
| CD68-PerCPCy5.5 | Biolegend | 137010 | FACS | 1:50 |
| PTPRC/CD45-eF450 | Thermo Fisher | 48-0451-82 | FACS | 1:100 |
| ITGAM/CD11b-APCeF780 | Thermo Fisher | 47-0112-82 | FACS | 1:100 |
| TREM2-AF647 | R&D Systems | FAB17291R | FACS | 1:100 |
| Cytokines and inflammasome: |  |  |  |  |
| TNFA-PE-Cy7 | Biolegend | 502929 | FACS | 1:100 |
| IL1B/IL-1β-PerCP-eF710 | NJTEN3, Invitrogen | 46-7114-82 | FACS | 1:100 |
| NLRP3-PE | Novus Biologicals | NBP2-12446PE | FACS | 1:100 |
| **Intracellular antibodies and markers:** |  |  |  |  |
| SQSTM1/P62-AF647 | Novus Biologicals | NBP1-42821AF647 | FACS | 1:50 |
| LC3B-FITC | Novus Biologicals | NB100-2220F | FACS | 1:100 |
| Lamp1-AF700 | Biolegend | 21627 | FACS | 1:100 |
| BODIPY-493/503 | Themro Fisher | D3922 | FACS | 1:5000 |
| GreenFluoroMyelin | Themro Fisher | F34651 | FACS | 1:500 |
| Galectin3/Mac2-PeCy7 | Biolegend | 125418 | FACS | 1:100 |
| Plin2 | Novus Biologicals | NB110-40877AF594 | FACS | 1:100 |

